# Evolution of sex-specific hormesis in a toxic niche specialist

**DOI:** 10.64898/2026.11.18.767557

**Authors:** Divyansh Mittal, Marie Messmer--Bachert, Michele Marconcini, Richard Benton

**Affiliations:** Center for Integrative Genomics, Faculty of Biology and Medicine, University of Lausanne, CH-1015, Lausanne Switzerland

## Abstract

Ecological specialists often exploit toxic niches that can exclude competitors, predators and pathogens. Whether such specialization has intrinsic fitness benefits remains unclear. We address this question through study of *Drosophila sechellia*, a specialist on noni fruit, which is toxic for other insects. A noni diet did not affect innate behaviors but increased female reproductive fitness, lifespan and stress resistance, and biased offspring sex ratios towards females. These traits were linked to female-specific lipid accumulation upon noni consumption. Strikingly, noni’s principal toxin, octanoic acid, reproduced these female-specific phenotypes, exhibiting a dose-dependent, biphasic –or hormetic–effect. The beneficial effects of octanoic acid require Alkbh7, a conserved mitochondrial protein involved in fatty acid metabolism. Finally, modelling showed fecundity and female bias can jointly maximize population growth. These results reveal how evolution of hormesis, to confer nutritious value upon a toxin, yields sexually-dimorphic fitness benefits, including an unusual uncoupling of reproductive fitness and lifespan.

## Introduction

Some of the most remarkable evolutionary adaptations of animals are exemplified by the specialization of species on toxic niches, such as crustaceans living by hydrothermal vents or in hypersaline environments (Chou et al., 2023; Santos et al., 2024; Tobler et al., 2016; Vetter et al., 1987; Zhang et al., 2024) and insects that feed on specific plants that are toxic for other animals (Beran and Petschenka, 2022; Dobler et al., 2012; Euw et al., 1967; Petschenka and Agrawal, 2016; Zhen et al., 2012). Exploitation of such ecological niches can prevent competition, predation and pathogenesis, but survival of the specialist requires its ability to deal with the toxic chemical(s) (Ali and Agrawal, 2012; Cornell and Hawkins, 2003; Futuyma and Moreno, 1988; Jaenike, 1990; Loxdale et al., 2011).

Several toxin tolerance and resistance strategies have been described (Tarvin et al., 2023). Chemicals can be detoxified by enzymes of the host or the gut microbiome, which contributes to pandas’ ability to avoid cyanide poisoning from bamboo (Zhu et al., 2018). Animals can exhibit increased excretion rate, as seen in desert woodrats when consuming creosote-bush toxins (Mangione et al., 2001), while other species cope with toxins through reduced physical activity, exemplified by koalas, which enables them to slowly process toxic eucalyptus chemicals (Johnson et al., 2018; Ryan et al., 2013). Some species harness consumed toxins for their own defense, such as poison dart frogs, which obtain toxic alkaloids from their diet of arthropods (Daly et al., 1994; Jeckel et al., 2026; Tarvin et al., 2024), or monarch butterflies, which sequester cardiac glycosides from their milkweed food source, rendering them unpalatable for predators (Agrawal, 2017; Agrawal et al., 2024; Jones et al., 2019). However, the dependency of ecological specialists on a single host for feeding and reproduction raises the question of whether they might also derive intrinsic fitness benefits from their toxic niche.

An interesting model to examine this question is *Drosophila sechellia* (Auer et al., 2021; Louis and David, 1986; Tsacas and Bächli, 1981), a fruit fly endemic to the Seychelles archipelago, which exhibits extreme specialization for the toxic “noni” fruit of the *Morinda citrifolia* shrub (Auer *et al*., 2021; Jones, 2005). Adaptation to this ecological niche has occurred in a relatively short time period, as ripe noni is highly toxic for close relatives of *D. sechellia*, notably *Drosophila simulans* and *Drosophila melanogaster* (Amlou et al., 1998b; Jones, 1998; 2001) (Figure 1A), as well as more divergent insects, including parasitoid wasps (Masuzzo et al., 2026; Salazar-Jaramillo and Wertheim, 2021). Specialization on noni is likely to reduce parasitization and interspecific competition for *D. sechellia* (Masuzzo *et al*., 2026; Salazar-Jaramillo and Wertheim, 2021), which is of particular importance given the low fecundity of females of this species compared to other drosophilids (Jones, 2004; R’Kha et al., 1997). Beyond the toxicity of noni for other species, early studies on *D. sechellia* noted that provision of noni increases egg-laying (R’Kha et al., 1991; R’Kha *et al*., 1997). However, whether noni has broader behavioral and life-history benefits have never been systematically examined, a question we set out to address in this work.

**Figure 1.**
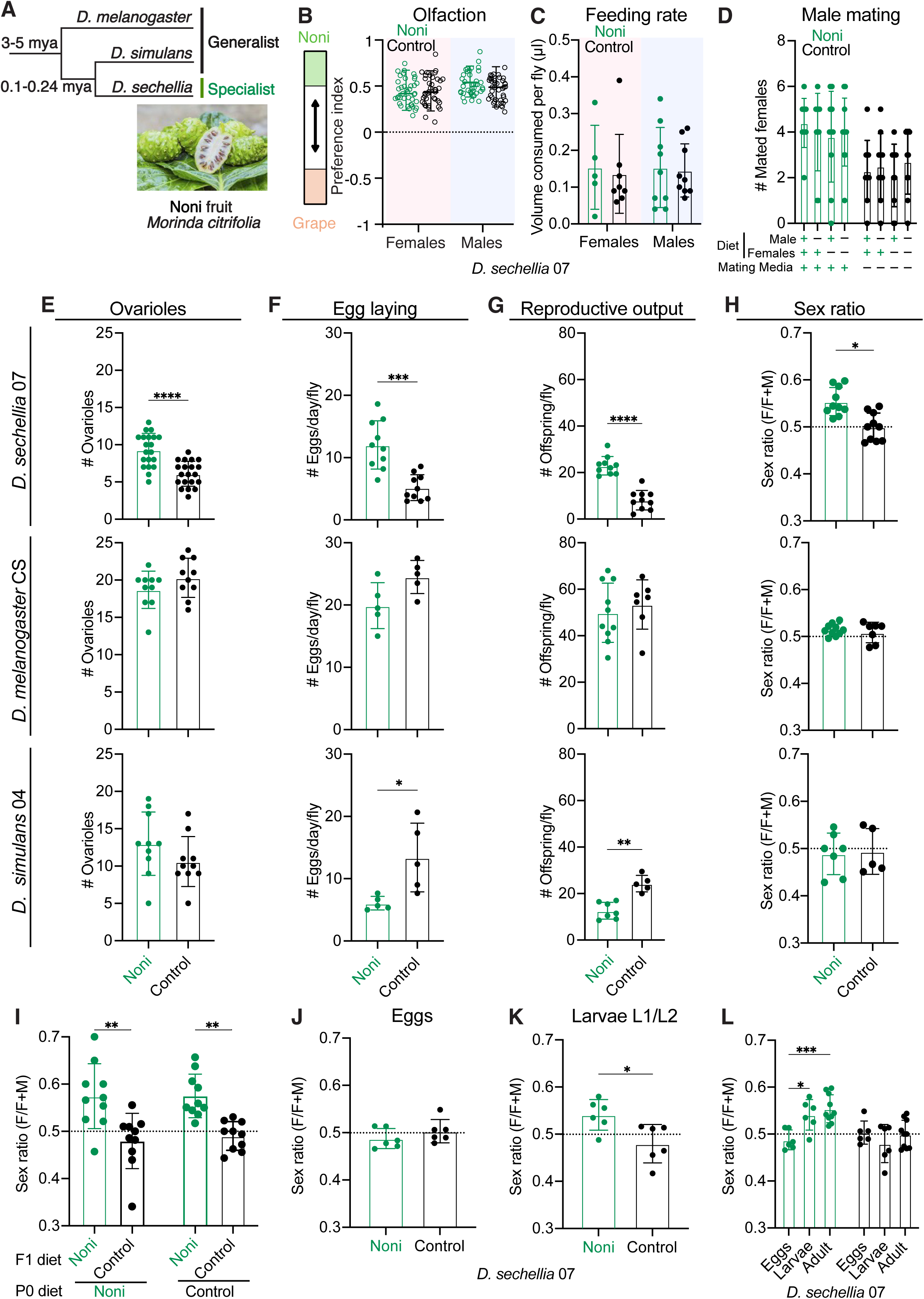
A noni diet enhances female reproductive fitness and shifts the sex ratio towards females in *D. sechellia*. (A) Phylogeny of the three species used in this study; times of last common ancestors are indicated in millions of years ago (Mya). (B) Preference index for noni juice versus grape juice for a two-choice olfactory assay using the planar arena for *D. sechellia* 07 reared on noni or control diet (*n* ≥ 40). (C) Feeding rate computed by measuring the volume of 2.5% sucrose solution consumed per fly using the CAFE assay (*n* ≥ 5 biological replicates of 10 flies each). (D) Mating capacity, quantified as the number of virgin females (out of 10 flies) successfully mated by a single male within a 24 h period (*n* = 10 replicates). (E-H) Female-specific reproductive traits and offspring sex ratio in specialist *D. sechellia* 07 (top row), and generalist *D. melanogaster* Canton-S (middle row) and *D. simulans* 04 (bottom row) fed noni or control diet: (E) mature ovariole number (*n* ≥ 10 females), (F) egg-laying rate (*n* ≥ 5 biological replicates of 10 females each), and (G) Reproductive output, quantified as the total number of adult offspring (females and males) emerging per replicate, divided by five to obtain offspring per female (*n* ≥ 7 biological replicates of 5 females each). (H) Offspring sex ratio (female offspring/total offspring) on noni and control diet (*n* ≥ 7 replicates). We note that noni juice is not toxic for generalists under the conditions used in these assays. (I) Effect of parental versus offspring developmental diet on offspring sex ratio of *D. sechellia* 07 *(n* ≥ 10 replicates). (J-K) Sex ratio computed by PCR-based sex identification in (J) eggs and (K) larvae produced by *D. sechellia* 07 fed a control or noni diet (*n* ≥ 6 replicates). (L) Sex ratio comparison across developmental stages (eggs, larvae and adult) and diets (noni and control). Data is same as H and J-K. Data in (B-C) were analyzed using two-way ANOVA with diet and sex as factors, followed by Šídák’s multiple-comparisons test to compare diets within each sex. Data in (D) were analyzed using a binomial generalized linear model (GLM), with mating medium, male diet and female diet as factors. Data in (E-H) and (J-K) were analyzed using two-sided Mann-Whitney U tests across diets. Data in (I) were analyzed using two-way ANOVA with parental diet and offspring developmental diet as factors, followed by Šídák’s multiple-comparisons test. Data in (L) were analyzed using two-way ANOVA with diet and offspring developmental stages as factors, followed by Šídák’s multiple-comparisons test for individual diets. Significance levels are indicated as \*\*\*\**P* < 0.0001, \*\*\**P* < 0.001, \*\**P* < 0.01, \**P* < 0.05; non-significant comparisons are not labeled.

## Results

### A noni diet does not alter attraction, feeding, locomotor or circadian behaviors in *D. sechellia*

We first asked whether a noni diet alters any innate behavioral responses of *D. sechellia*. Flies reared on a noni juice or control diet (see Methods) showed similar attraction to noni odors in a two-choice olfactory assay, confirmed in both sexes and against two competing stimuli (Figure 1B and Figure S1A), extending previous observations (Auer et al., 2020). Feeding rates on sucrose solution or noni juice, measured by the Capillary Feeder (CAFE) assay, were likewise indistinguishable in noni- and control-diet cultured flies of either sex (Figure 1C and Figure S1B). Similarly, circadian locomotor activity and sleep, measured by the *Drosophila* Activity Monitor, were not influenced by the flies’ diet (Figure S1C-F). The absence of a dramatic effect of a noni diet on these innate behaviors is consistent with the limited effect of noni consumption on the cellular composition of, and gene expression patterns in, the *D. sechellia* brain (Lee et al., 2025).

### A noni diet enhances female, but not male, reproductive fitness in *D. sechellia*

We next examined the effect of noni on the reproductive fitness of *D. sechellia*. Male mating success increased when the mating assay itself was conducted on noni compared to control medium (Figure 1D), indicating that the acute presence of noni promotes mating, reminiscent of the positive effect of food on male courtship in other drosophilid species (Brand et al., 2026; Grosjean et al., 2011). However, the rearing diet of the tested flies did not affect mating frequency in either sex (Figure 1D), nor was there an effect of male diet on the number of offspring produced per inseminated female (Figure S2A).

By contrast, in females, a noni diet led to a higher number of mature ovarioles (Figure 1E), similar to previous observations (Lavista-Llanos et al., 2014). Females shifted from control to noni diet as adults showed an increase in mature ovarioles, whereas those shifted from noni to control diet showed a decrease (Figure S2B-C), indicating that ovariole maturation is most sensitive to adult diet. Consistent with these observations, noni-reared females laid significantly more eggs over a three-day period (Figure 1F). These eggs also exhibited higher viability (Figure S2D) and altered timing of eclosion (Figure S2F-G); consequently, females maintained on noni throughout their life produced substantially more offspring than controls (Figure 1G and Figure S2E). The reproductive age of *D. sechellia* was not, however, noticeably extended by a noni diet (Figure S2F). In contrast to its effect on *D. sechellia*, noni did not enhance female reproductive output in generalist drosophilid species (Figure 1E-G and Figure S2G-H), but rather decreased it, likely reflecting the unfavorable nature of noni.

We validated the beneficial effect of noni on *D. sechellia*’s female reproduction in several ways. First, rearing flies on an isocaloric grape juice diet did not lead to an increase in female reproductive output (Figure S2I). Second, rearing flies on ripe noni fruit produced a similar effect as the noni juice diet (Figure S2I). Third, we confirmed the beneficial effects of the noni diet in an independent laboratory strain of *D. sechellia* (Figure S2J-M) and in recently wild-caught *D. sechellia* strains (Shahandeh et al., 2026) (Figure S2N-O). Together, these results show that noni enhances multiple aspects of female reproductive fitness specifically in *D. sechellia*.

### Female-biased sex ratio in noni-reared *D. sechellia*

In the course of these experiments, we serendipitously observed that offspring of noni-reared, but not control-reared, *D. sechellia* exhibited a female-biased sex ratio (Figure 1H). This phenomenon was observed in other laboratory and wild-caught strains (Figure S3A-B), but not generalist species (Figure 1H and Figure S3C-D). The shift was also observed when flies were reared on noni fruit, but not on an isocaloric grape juice diet (Figure S3E-F), nor when *D. sechellia* was cultured at higher or lower temperatures (Figure S3G) (which affects sex determination in other taxa (Ferguson and Joanen, 1982; Pezaro et al., 2017; Quinn et al., 2007; Warner and Shine, 2008)).

We investigated whether noni impacts sex ratio through an influence on the parental generation (i.e., on their gametes) and/or arises during offspring development, by rearing on the noni diet and transferring them to the control diet as adults for egg laying and development, or vice versa. Offspring developing on noni exhibited a female-biased sex ratio regardless of the parents’ original diet (Figure 1I), indicating that the effect arises after fertilization rather than through primary sex allocation.

To determine when during development this bias emerges, we used PCR-based sexing of eggs and larvae. Sex ratio was balanced at the egg stage but became female-biased on noni by the first/second larval instar stage (Figure 1J-K). A similar result was obtained using an RFP-marked X-chromosome to determine the sex of offspring (Figure S3H-I). On a noni diet, the larval offspring sex ratio shift was similar to adult sex ratio shift (Figure 1L). These findings identify early larval development as the stage when noni transforms a sex-ratio balanced cohort into a female-biased population; the potential benefit of this phenomenon is considered below (Figure 6).

### Noni selectively improves female lifespan

The reproductive benefits of noni raised the question of whether females gain reproductive fitness at the cost of longevity, a trade-off observed in many other species (Flatt, 2011; 2020; Fowler and Partridge, 1989; Harshman and Zera, 2007; Lee et al., 2008; Travers et al., 2015; Zera and Harshman, 2001). Unexpectedly, noni-reared females exhibited a significant increase in lifespan compared to control-reared females, whereas male lifespan was unaffected (Figure 2A). This female-specific effect was also observed when flies were reared on noni fruit, but not on a non-noni, isocaloric diet (Figure S4A-D). To determine whether adult diet is sufficient to modulate lifespan, flies were switched from their developmental diet to the alternative diet upon adulthood. Females shifted from control to noni diet as adults exhibited increased lifespan compared to those shifted from noni to control diet (Figure 2B). Again, no such differences were observed in males (Figure 2B).

**Figure 2.**
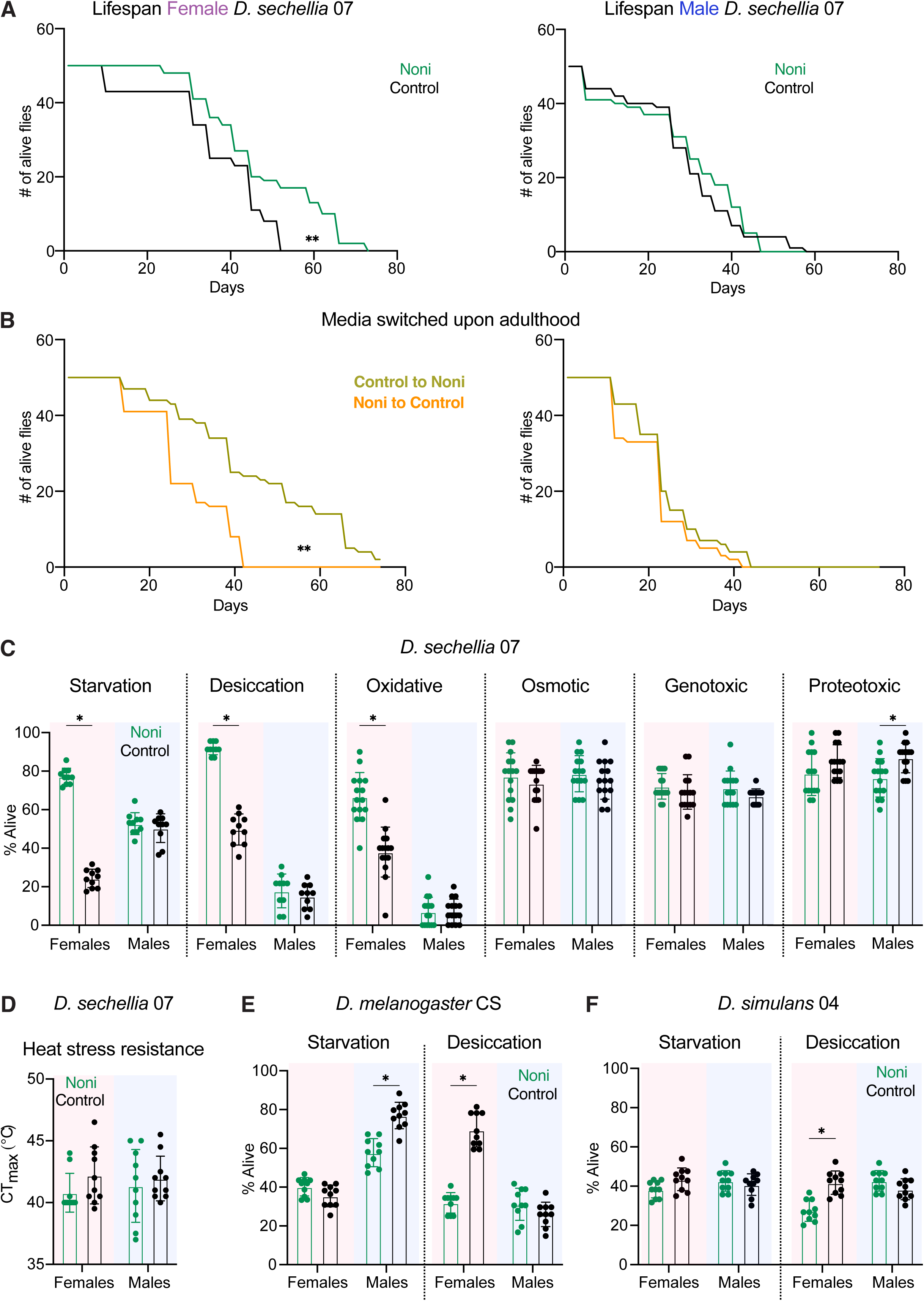
Noni diet improves lifespan and stress resistance traits in female, but not male, *D. sechellia*. (A) Lifespan of females (left) and males (right) of *D. sechellia* 07 maintained on noni or control diets (*n* = 50 flies). (B) Lifespan of females (left) and males (right) of *D. sechellia* 07 developed on control diet and switched to noni diet upon eclosion and maintained until death (or vice versa) (*n* = 50 flies). (C) Stress resistance of *D. sechellia* 07 maintained on noni or control diet. Starvation, desiccation, oxidative stress following 18.5 mM paraquat exposure, genotoxic stress following 15 mM CuSO_4_ exposure, osmotic stress following 400 mM NaCl exposure, and proteotoxic stress resistance following 5 mM CdCl_2_ exposure were tested (*n* ≥ 10 replicates). (D) Heat stress resistance (Critical Thermal Maximum, CT_max_) of *D. sechellia* 07 reared on a noni or control diet (*n* ≥ 10). (E-F) Starvation and desiccation resistance of generalist (E) *D. melanogaster* Canton-S and (F) *D. simulans* 04 maintained on noni or control diet (*n* ≥ 10 replicates). Survival differences in (A-B) were assessed using the log-rank (Mantel–Cox) test (*n* = 50 flies per sex and diet). Data in (C-F) were analyzed using two-way ANOVA with diet and sex as factors, followed by Šídák’s multiple-comparisons test to compare diets within each sex. Significance levels are indicated as **** *P* < 0.0001, *** *P* < 0.001, ** *P* < 0.01, * *P* < 0.05; non-significant comparisons are not labeled.

Because noni enhances both female reproductive fitness and lifespan, we asked whether the longevity benefit depends on reproduction itself. Surprisingly, control-reared virgin females (and males) had dramatically decreased lifespan (Figure S4E-F), a phenotype that is distinct from other drosophilids (Aigaki and Ohba, 1984; Fowler and Partridge, 1989; Kim et al., 2010; Taylor et al., 2008; Travers et al., 2015). This effect reinforces the evidence for decoupling of reproduction and lifespan in *D. sechellia*. Although the noni diet increased virgins’ lifespan, the effect was small and non-sexually dimorphic (Figure S4E-F). Together, these findings indicate that noni enhances female lifespan without an accompanying reduction in reproductive performance, suggesting that the relationship between reproduction and longevity is fundamentally altered in this specialist.

### Noni selectively improves female stress resistance

We next investigated whether noni has broader benefits by assessing resistance to a range of environmental stressors. Noni-reared females exhibited increased resistance to starvation, desiccation and oxidative stress, but not osmotic, genotoxic, proteotoxic or heat stress (Figure 2C-D). The improved starvation resistance is unlikely to result from altered feeding rate (Figure S5A), metabolic rate (Figure S5B) nor behavioral energy conservation as noni-reared females exhibited similar starvation-induced locomotor responses and sleep patterns compared to flies grown on the control diet (Figure S5C-G). In *D. sechellia* males (Figure 2C) or generalist drosophilid species (Figure 2E-F), noni did not produce beneficial effects on stress resistance; rather, in certain cases, it reduced the ability of flies to survive the stressor.

### Noni diet induces female-specific lipid accumulation and metabolic remodeling

To explore the physiological changes that might explain the female-specific fitness benefits, we asked how noni influences body composition and metabolism of *D. sechellia*. Rearing on noni increased both wet and dry body weight in females but not males (Figure 3A-B), an effect not seen in generalist drosophilids (Figure S6A-B). Whole-body, untargeted metabolomic profiling of *D. sechellia* revealed a considerably stronger noni-dependent diet response in females compared to males: of the 313 total metabolites detected (201 using multi-pathway targeted analysis and 112 using untargeted metabolomic analysis), females exhibited 22 differentially-abundant metabolites (5 and 17 at higher and lower levels, respectively, in noni-fed flies), compared to only 3 in males (all at higher levels, which were also found in females) (Figure 3C and Figure S6C-E). The more abundant metabolites shared between sexes were gelseminic acid (scopoletin), malonate and trigonelline; as noni is rich in these compounds (Almeida et al., 2019; Liu et al., 2024; Pino et al., 2009), we suspect the higher levels in noni-fed *D. sechellia* reflect their direct dietary origin (as is probably also the case for rose oxide, which is more abundant only in females). The female-specific, lower abundance chemicals were matched sparsely to multiple pathways, including amino acid metabolism and oxidative energy production (Figure S6F).

**Figure 3.**
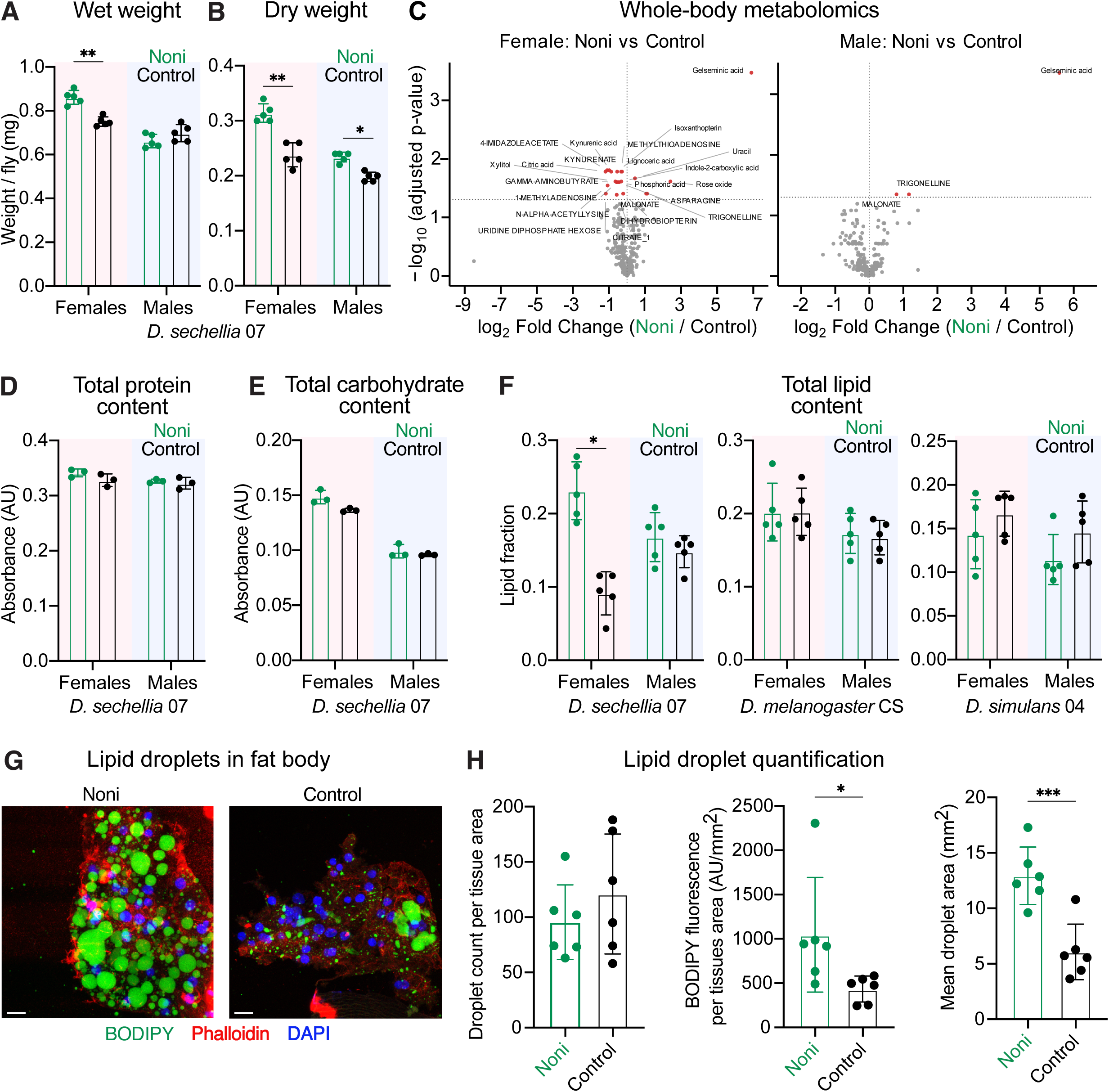
Noni diet induces sex-specific metabolomic remodeling and lipid accumulation in *D. sechellia*. (A) Wet body weight of *D. sechellia* 07 females and males on noni or control diet (*n* = 5 biological replicates of 10 flies each). (B) Dry body weight of *D. sechellia* 07 females and males on noni or control diet (*n* = 5 biological replicates of 10 flies each). (C) Metabolomics volcano plot for *D. sechellia* 07 females (left) and males (right) on noni versus control diet. Dotted lines denote FDR 0.05 thresholds on y-axis and log_2_FC = 0 on x-axis. Significantly differentially abundant metabolites are highlighted in red. (*n* = 5 samples per sex per diet treatment). (D) Total protein content of *D. sechellia* 07 females and males on noni or control diet (*n* = 3 biological replicates). (E) Total carbohydrate (glucose) content of *D. sechellia* 07 females and males on noni or control diet (*n* = 3 biological replicates). (F) Total lipid content of *D. sechellia* 07 (left), *D. melanogaster* CS (center) and *D. simulans* 04 (right) females and males on noni or control diet (*n* = 5 biological replicates). (G) Representation images of lipid droplets in fat body of female *D. sechellia* 07 on noni or control diet. BODIPY (green), Phalloidin (red) and DAPI (blue). Scale bars, 10 μm. (H) BODIPY Fluorescence was used to quantify lipid droplets in fat body: (left) droplet count normalized to tissue area, (middle) background corrected BODIPY fluorescence intensity, and (right) mean droplet area and summarized per biological replicate (*n* = 6 biological replicates). Data in (A-F) were analyzed using two-way ANOVA with diet and sex as factors, followed by Šídák’s multiple-comparisons test to compare diets within each sex. Data in (H) were analyzed using two-sided Mann–Whitney U tests. Significance levels are indicated as **** *P* < 0.0001, *** *P* < 0.001, ** *P* < 0.01, * *P* < 0.05; non-significant comparisons are not labeled.

As this analysis did not highlight a single metabolic pathway that was influenced by noni, we performed a broader analysis of the macronutrient composition of flies. While total protein or carbohydrate (glucose) contents of animals were not affected by diet, lipid content increased significantly in females (Figure 3D-F). This result implied that elevated lipid stores could account for the increase in body weight. No comparable noni diet-dependent changes in body weight or macronutrient composition were observed in either sex of generalist species (Figure 3F and Figure S6G-H).

We directly visualized lipid accumulation *in vivo* of female *D. sechellia* by BODIPY staining of the fat body, the main energy storage organ of insects (analogous to the mammalian adipose tissue and liver (Li et al., 2019)). The number of lipid droplet was not different between noni-and control-reared females, we observed increased lipid-droplet fluorescence intensity and size (Figure 3G-H). These results indicate increased lipid storage in the fat body, likely primarily through enlargement of existing lipid droplets. These may provide females with an increased energy reserve to support reproductive and somatic functions (see Discussion).

### The main noni toxin, octanoic acid, improves female reproductive capacity and health benefits

We next investigated which component of noni is responsible for the beneficial effects on *D. sechellia*. Varying the proportion of noni in the diet led to dose-dependent changes in egg laying, reproductive output, sex ratio and starvation resistance (Figure S7A-D), suggesting that the key chemical(s) is a macronutrient component of noni. We screened several such noni constituents across a range of concentrations – including fatty acids, sugars, and unique components like scopoletin and L-DOPA – for their ability to stimulate egg laying compared to a matched control diet (Figure 4A and Figure S7E). The sole compound capable of reproducing this response was octanoic acid, a medium-chain fatty acid abundant in noni. This unique property of octanoic acid was further highlighted by testing related short- and medium-chain fatty acids diets as well as other eight-carbon compounds: none of these stimulated egg-laying at any concentration (Figure 4A and Figure S7E-F).

**Figure 4.**
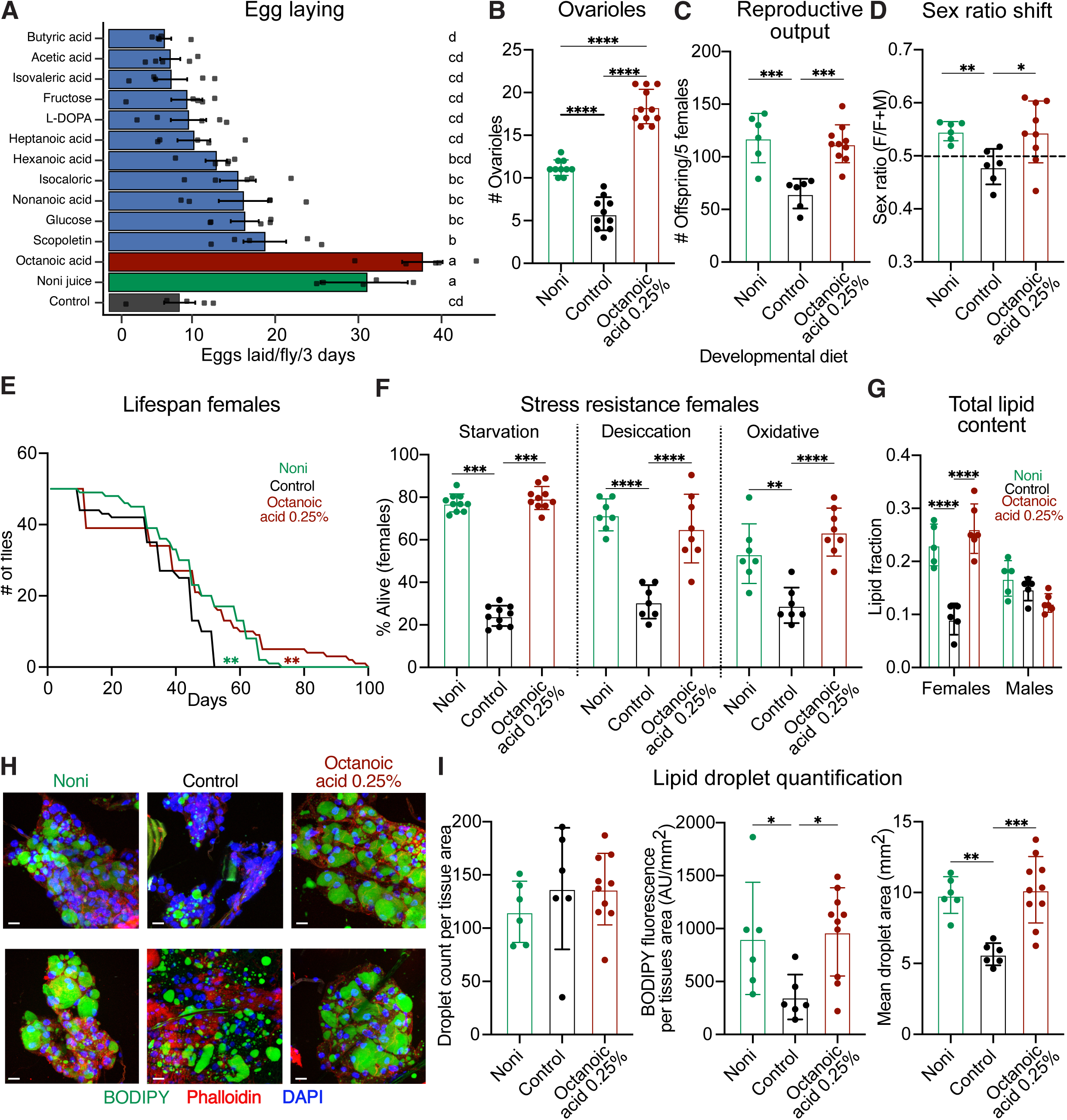
Octanoic acid in noni is a driver of sex-specific metabolic evolution and sex ratio shifts at larval stages. (A) Screen of compounds found in noni fruit for egg-laying (*n* ≥ 5). For each compound (mixed in control diet), the concentration eliciting the highest egg-laying response is shown; data for all five concentrations are presented in Figure S7E. (B-D) Effect of octanoic acid diet (0.25% v/v) on (B) mature ovariole number, (C) reproductive output, (D) offspring sex ratio of *D. sechellia 07* (*n* ≥ 6). (E) Lifespan of females *D. sechellia* 07 maintained on noni, control or octanoic acid (0.25% v/v) diet (*n* = 50 flies). (F) Stress resistance of *D. sechellia* 07 females maintained on noni, control or octanoic acid (0.25% v/v) diet. Starvation, desiccation, oxidative stress following 18.5 mM paraquat exposure (*n* ≥ 10 replicates). (G) Total lipid content of *D. sechellia* 07 females and males on noni, control or octanoic acid (0.25% v/v) diet (*n* = 5 biological replicates). (H) Representation images of lipid droplets in fat body of female *D. sechellia* 07 on noni, control or octanoic acid (0.25% v/v) diet. BODIPY (green), Phalloidin (red) and DAPI (blue). Scale bars, 10 μm. (I) BODIPY Fluorescence was used to quantify lipid droplets in fat body: (left) droplet count normalized to tissue area, (middle) background corrected BODIPY fluorescence intensity, and (right) mean droplet area and summarized per biological replicate (*n* ≥ 6 biological replicates). Data in (A-D), (F) and (I) were analyzed using one-way ANOVA, followed by Tukey’s multiple-comparisons test. Survival differences in (E) were assessed using the log-rank (Mantel–Cox) test. Data in (G) were analyzed using two-way ANOVA with diet and sex as factors, followed by Tukey’s multiple-comparisons test to compare diets within each sex. Significance levels are indicated as **** *P* < 0.0001, *** *P* < 0.001, ** *P* < 0.01, * *P* < 0.05; non-significant comparisons are not labeled.

In addition to egg-laying, octanoic acid increased ovariole maturation (Figure 4B), reproductive output (Figure 4C), as well as reproducing the noni-dependent female-biased shift in sex ratio at the larval stage (Figure 4D and Figure S8A-B). Furthermore, octanoic acid improved female lifespan (Figure 4E), resistance to starvation, desiccation and oxidative stress (Figure 4F). No such octanoic acid diet-dependent improvement was observed for males (Figure S8C-D). *D. melanogaster* did not lay eggs and died on the matching octanoic acid diet (Figure S8E). Lastly, the octanoic acid diet led to increases in both wet and dry weights (Figure S8F) as well as total lipid content (Figure 4G) and increased lipid accumulation in the fat body (Figure 4H-I). Together, these results highlight octanoic acid as a single component of noni that can recapitulate all the key effects of the noni diet.

### Octanoic acid exhibits species-specific hormesis

The beneficial effects of octanoic acid for *D. sechellia* were unexpected as this chemical is well-established to be the principal toxic component of noni for other insect species (Amlou *et al*., 1998b; Jones, 1998; 2001; Masuzzo *et al*., 2026; Salazar-Jaramillo and Wertheim, 2021). To understand this apparent paradox, we examined the dose-dependent effects of octanoic acid on both viability and egg-laying rate of *D. sechellia*. Females tolerate extremely well a wide range of octanoic acid concentrations, but eventually succumb to its toxic effects (Figure 5A). Males exhibit a similar tolerance profile (Figure 5B). Within the range at which *D. sechellia* exhibits full viability, egg-laying rate exhibited a narrow peak at 0.25-0.5% (v/v), where it lays ∼7-fold more eggs than animals on a control diet (Figure 5C; see also Figure S7E). (This sensitive dose-dependency of octanoic acid might explain the inconsistencies in previous observations of the effect of octanoic acid on oviposition preference and/or rate (Alvarez-Ocana et al., 2023; Amlou et al., 1998a; Higa and Fuyama, 1993; Legal et al., 1999; Markow et al., 2009; Matsuo et al., 2007)). These results highlight a dose-dependent, biphasic – or hormetic (Calabrese and Baldwin, 2003; Rix and Cutler, 2022) – mode-of-action of octanoic acid, in which low levels have beneficial effect, while high levels are harmful. The optimal beneficial concentration of octanoic acid corresponds remarkably well with the levels found in noni fruit (3.06 g/kg fresh fruit (Pino *et al*., 2009), equivalent to ∼0.34% v/v in our medium, after accounting for base medium and octanoic acid densities). This close match suggests that *D. sechellia*’s capacity to use octanoic acid for benefit is a specific adaptation to its ecological niche.

**Figure 5.**
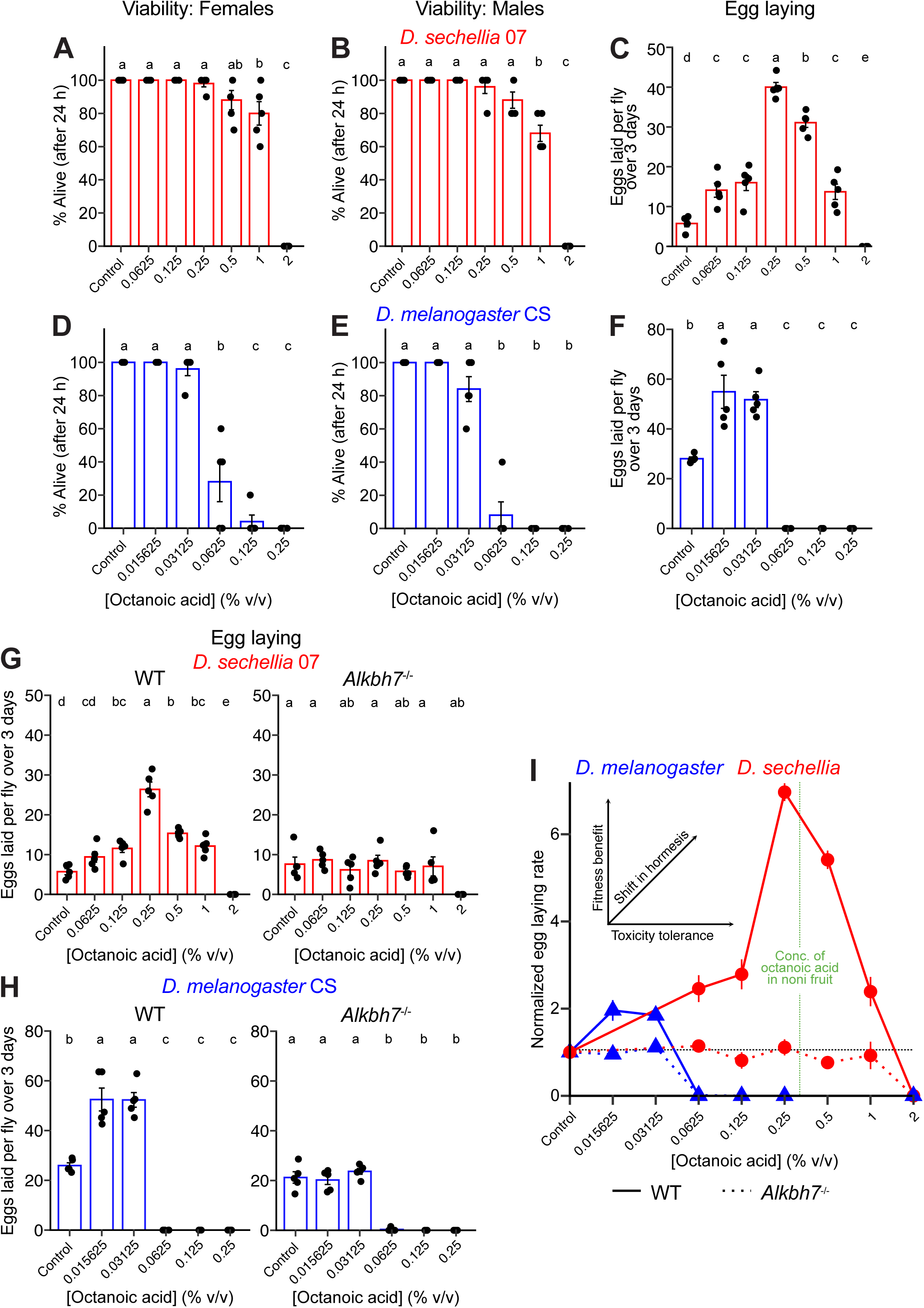
Octanoic acid exhibits species-specific hormesis. (A-B) Percentage survival of *D. sechellia* 07 (A) females and (B) males after 24 h exposure to control medium supplemented with the indicated concentrations of octanoic acid (*n* = 5 replicates of 10 flies each). (C) Egg-laying response of female *D. sechellia* 07 to control medium supplemented with the indicated concentrations of octanoic acid (*n* = 5 replicates of 10 flies each). (D-E) Percentage survival of *D. melanogaster* CS (D) females and (E) males after 24 h exposure to nutrient-poor medium (0.25×) supplemented with the indicated concentrations of octanoic acid (*n* = 5 replicates of 5 flies each). (F) Egg-laying response of female *D. melanogaster* CS to nutrient-poor medium (0.25×) supplemented with the indicated concentrations of octanoic acid (*n* = 5 replicates of 5 flies each). In normal control medium (1×) (Figure S8E), the egg-laying rate of *D. melanogaster* was already very high and not affected by octanoic acid supplementation. (G) Egg-laying response of *D. sechellia* 07 and *Alkbh7* mutant *(Alkbh7^RFP^*, in the 07 background) to control medium supplemented with the indicated concentrations of octanoic acid (*n* = 5 replicates of 10 flies each). (H) Egg-laying response of *D. melanogaster* CS and *Alkbh7* mutant (*Alkbh7^1^*, in the CS background) to nutrient-poor medium (0.25×) supplemented with the indicated concentrations of octanoic acid (*n* = 5 replicates of 5 flies each). (I) Hormetic egg-laying responses of *D. sechellia* (07 and *Alkbh7* mutant) and *D. melanogaster* (CS and *Alkbh7* mutant), normalized to the respective control response for each species (horizontal black dashed line). The vertical green dashed line indicates the estimated equivalent of the octanoic acid concentration reported in fresh noni fruit (3.06 g kg^-1^ (Pino *et al*., 2009), equivalent to approximately 0.34% v/v in our medium;). Data in (A-H) were analyzed using one-way ANOVA, followed by Tukey’s multiple-comparisons test.

We wondered whether octanoic acid hormesis is a unique phenomenon for *D. sechellia* or is detectable in other species. While *D. melanogaster* females are much less tolerant of octanoic acid, with only ∼30% survival at the lowest concentration tested in our dose-response analysis of *D. sechellia* (0.0625% v/v), further dilution of this chemical does permit survival of *D. melanogaster* (Figure 5D-E). Notably, at these concentrations, *D. melanogaster* exhibits a modest (<2-fold) increase in egg-laying rate compared to flies on a control medium (Figure 5F). These observations suggest that octanoic acid also exhibits hormesis for *D. melanogaster*, albeit with distinct concentration thresholds and magnitudes of effect.

How octanoic acid exerts dose-dependent negative and positive effects is likely to be complex: despite decades of research, the mechanism underlying its toxicity – and *D. sechellia*’s high resistance – is still unclear (see Discussion), but we hypothesized that the beneficial effects of octanoic acid depend upon metabolic utilization of this medium-chain fatty acid. To explore this idea, we took advantage of *D. sechellia* mutants lacking *Alkbh7*, which encodes an evolutionarily-conserved mitochondrial protein required, in mammals, for fatty acid metabolism, likely through regulation of protein translation in mitochondria (Solberg et al., 2013; Zhang et al., 2021). In drosophilids, *Alkbh7* previously-emerged from artificial selection screens as a gene contributing weakly to tolerance of octanoic acid (Marconcini et al., 2026a). We asked whether loss of *Alkbh7* impacts the beneficial effects observed upon exposure to non-lethal doses of this fatty acid. Strikingly, *D. sechellia Alkbh7* null mutants completely lacked the octanoic acid-induced increase in egg-laying (Figure 5G). A similar loss of the beneficial effects of octanoic acid on egg-laying were seen in *D. melanogaster Alkbh7* mutants (Figure 5H). These observations support a model in which mitochondrial-based metabolism of octanoic acid (or a derivative), through an as-yet unknown pathway, underlies the beneficial phase of hormesis (see Discussion; Figure 5I).

### A demographic model reveals fecundity and female bias jointly shape population growth

Testing if and how *D. sechellia* populations might be affected by the female-specific fecundity benefit, the female-skewed sex ratio and survival benefit (representing lifespan and stress resistance) associated with noni and octanoic acid is difficult in the wild. We therefore developed a phenomenological, stochastic, two-sex population model (Figure 6A). We first established a control scenario in which the starting population was arbitrarily set at 200 animals (100 of each sex), the carrying capacity arbitrarily set at 1000. Baseline birth and death rates were adjusted to maintain an approximately stable population. All simulations began with this population size, providing a common reference against which to assess the effects of changing different traits.

**Figure 6.**
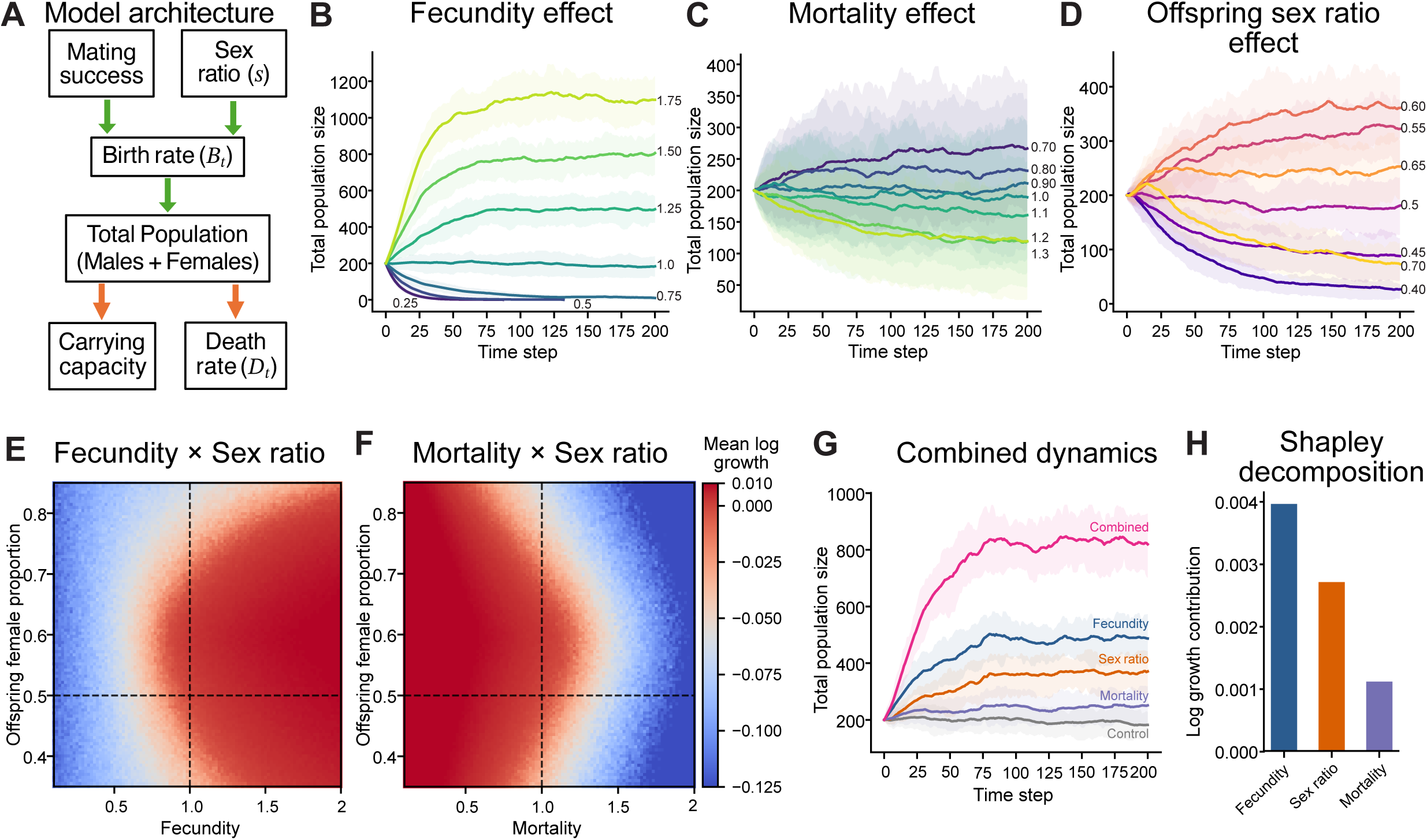
Female-biased fitness benefits can reshape population dynamics. (A) Schematic of the stochastic two-sex population model. (B) Simulated population trajectories across fecundity multipliers from 0.25× to 1.75× baseline fecundity. (C) Simulated population trajectories across mortality multipliers from 0.70× to 1.30× baseline mortality. (D) Simulated population trajectories across offspring sex ratios, from 40% to 70% female. In (B-D), lines represent mean population size across 100 stochastic simulations per condition, and shaded regions indicate the interquartile range (25th–75th percentiles). (E) Heatmap of mean log population growth per time step across combinations of fecundity multiplier and female fraction among offspring. The 100 × 100 parameter grid spans fecundity multipliers from 0.1× to 2.0× and female fractions from 0.35 to 0.85; each grid point represents the mean of 30 stochastic simulations. (F) Heatmap of mean log population growth per time step across combinations of mortality multiplier and female fraction among offspring. The 100 × 100 parameter grid spans mortality multipliers from 0.1× to 2.0× and female fractions from 0.35 to 0.85; each grid point represents the mean of 30 stochastic simulations. (G) Simulated population trajectories under control, fecundity-only, sex-ratio-only, mortality-only, and combined treatment scenarios. The combined treatment assumes a 1.25-fold increase in fecundity, a female offspring fraction of 0.60, and a mortality multiplier of 0.95. Lines represent mean population size across 60 stochastic simulations per condition, and shaded regions indicate the interquartile range (25th–75th percentiles). (H) Shapley decomposition of the contributions of fecundity, offspring sex ratio, and mortality to the difference in population growth between control and combined treatment scenarios, calculated using 200 stochastic simulations per component combination.

Increasing fecundity consistently promoted population growth, as expected, whereas low fecundity led to decline (Figure 6B). Mortality also had the expected effects on population growth, although these were more limited (Figure 6C). Importantly, offspring sex ratio had a nonlinear effect on population growth: populations stabilized at the largest size at a moderate female bias (∼55-60% female), whereas stronger bias diminished this benefit as males became limiting for fertilization (Figure 6D). Notably, the optimal sex ratio for population growth in these simulations matched that observed in our experiments (55-60% (Figure 1H)). Population growth was comparatively insensitive to all other tested factors, for example, environmental noise and initial female proportion (Figure S9). One exception was male fertilization capacity: where this is too low, insufficient females are fertilized, leading to population decline (Figure S9), paralleling the consequence of a strongly-female biased sex ratio (Figure 6D).

We next more comprehensively tested the effects of fecundity and offspring sex ratio with 10,000 combinations of these parameters. Higher fecundity always increased growth, and growth peaked at an offspring sex ratio of ∼60% female whatever the fecundity, rather than at the strongest female bias (Figure 6E). A similar analysis with mortality and sex ratio also pointed toward stable population growth with moderate female-biased sex ratio (Figure 6F).

We next asked how the three traits act together. Applying each trait alone increased population size relative to control; combining all three increased it more than four-fold (Figure 6G). To apportion this combined gain among the three traits, we used a Shapley decomposition, which averages each trait’s added effect across all orders in which the traits could be introduced. Fecundity was the largest contributor (∼50% of the gain in growth), followed by offspring sex ratio (∼35%) and reduced mortality (∼15%) (Figure 6H). Together, these simulations suggest that noni-associated fitness benefits extend beyond individual performance, with increased fecundity and a moderately female-biased offspring sex ratio most strongly favoring population growth and persistence.

## Discussion

Specialization on toxic ecological niches has largely been interpreted as a strategy that enables a species to alleviate competition for resources and protect against potential predators or pathogens. As such, studies of the underlying adaptations have focused on sensory mechanisms to locate and identify such niches and defense mechanisms to resist or tolerate the toxic chemicals. Here, we have revealed how an extreme specialist, the fly *D. sechellia*, has gained widespread fitness benefits from its toxic noni niche, spanning reproduction, lifespan and stress resistance. These benefits highlight an unexpected absence in *D. sechellia* of the trade-off between lifespan and reproductive output that is seen in many other animals (Flatt, 2011; Lemaître et al., 2015; Promislow and Harvey, 1990; Wilder et al., 2013), suggesting that this drosophilid accesses additional energy resources that render this trade-off unnecessary. Strikingly, we discovered that the principal noni toxin, octanoic acid, is also the main inducer of these fitness increases. The beneficial and harmful effects of octanoic acid are dose-dependent, exemplifying the process of hormesis (Calabrese and Baldwin, 2003; Rix and Cutler, 2022). Our observation that octanoic acid has a measurable, albeit small, hormetic effect in *D. melanogaster* indicates that this phenomenon existed already in the ancestor of *D. melanogaster* and *D. sechellia*. The adaptation of *D. sechellia* to its niche has therefore involved the evolution of hormesis, in which the threshold and magnitude of its positive and negative effects have changed (Figure 5I). Notably, specialization has repositioned octanoic acid’s ecologically-relevant dose (in noni fruit) within a beneficial range for *D. sechellia*, rather than simply raising the threshold at which it becomes harmful (Figure 5I).

Determining the mechanistic basis of octanoic acid’s hormetic effect remains an important future challenge. Diverse genetic and genomic analyses of octanoic acid susceptibility/resistance in various drosophilid species have pointed to multiple possible defense mechanisms – including increased cuticular barrier function, detoxification and altered metabolism (Amlou *et al*., 1998b; Amlou et al., 1997; Ando et al., 2019; Huang and Erezyilmaz, 2015; Jones, 1998; Lanno et al., 2017; López et al., 2017; Marconcini *et al*., 2026a; Marconcini et al., 2026b; Sun et al., 2024) – although these insights have not revealed the toxin mode-of-action *per se*. The mechanism by which *D. sechellia* derives benefit from octanoic acid is also unclear, but two observations are informative. First, the accumulation of lipid droplets in the fat body in flies on a noni or octanoic acid diet raises the possibility that octanoic acid is itself incorporated into these droplets through lipogenesis, for example, by elongation of this fatty acid and incorporation into triglycerides that concentrate in the fat body. Testing this hypothesis will require isotope tracing studies, but it is appealing to account for both a detoxification route and the subsequent beneficial effects we have observed. In *D. melanogaster* (and presumably other drosophilids), lipid droplets are key energy reserves that have been linked to starvation and oxidative stress resistance, lifespan increases and female fertility (Almeida-Oliveira et al., 2021; Arrese and Soulages, 2010; Bailey et al., 2015; Ballard et al., 2008; Briegel et al., 2002; Kalra and Parkash, 2014; Rehman and Varghese, 2021; Sreejith et al., 2024; Yang et al., 2024). Second, our analysis of mutants lacking *Alkbh7* provide a genetic link, in both *D. sechellia* and *D. melanogaster*, between fatty acid metabolism and the reproductive benefits of a non-lethal dietary supplement of octanoic acid. Future dissection of fatty acid metabolism under different dietary regimes and across species will be necessary to define the mechanistic links between these phenomena.

It seems possible, if not probable, that octanoic acid toxicity, resistance and benefit are mechanistically intertwined; by increasing use of octanoic acid for benefit, *D. sechellia* avoids the negative effects of this chemical present at high levels in its niche. The *Alkbh7* phenotype provides some support for this: in addition to loss of octanoic acid-induced egg-laying, *D. sechellia* mutants also have reduced tolerance of high concentrations of octanoic acid over longer timescales (Marconcini *et al*., 2026a). A second hint of the coupling of these processes comes from analysis of the fat body-expressed CD36-related transporter Bez: in *D. melanogaster*, this protein both mediates lipid export from this organ to support ovariole maturation (Carrera et al., 2024) and contributes to tolerance of octanoic acid exposure (Marconcini *et al*., 2026b). Whatever the precise cellular, molecular and/or metabolic mechanisms involved, we speculate that *D. sechellia*’s ability to funnel octanoic acid towards beneficial effects is more efficient than that of generalist drosophilids, resulting in the fundamental differences in how these species respond to exposure to this chemical. In this context, we note that *Alkbh7* is expressed at much higher levels in *D. sechellia* compared to *D. melanogaster* (Marconcini *et al*., 2026a), pointing towards a potential mechanism underlying the species differences in the hormetic effect of octanoic acid.

Finally, our results reveal a cryptic physiological sexual dimorphism in *D. sechellia*: females derive greater life-history and fitness benefits from noni than males. Adaptation to novel ecological niches is typically considered at the level of the species (Connallon et al., 2018; De Lisle, 2019), but our results show that adaptation can selectively reshape the biology of only one sex, emphasizing the importance of studying both sexes when considering evolutionary change. A notable consequence of trait differences at the level of individuals is a female-biased secondary sex ratio, which presumably reflects increased mortality of males and/or increased viability of females during development on noni. What the precise cause, this phenomenon provides, to our knowledge, an unprecedented example where a specialized food source directly skews the sex ratio of a population. While population sex ratio is the outcome rather than the trait under selection (which acts at the level of an individual), our models suggest that the demographic impact of a female-skewed ratio might help counter-balance *D. sechellia*’s lower female fecundity to stabilize population size. However, whether noni-dependent survival differences generate comparable sex-ratio shifts in wild populations, and what consequences these shifts have for natural population dynamics, remain to be tested.

“*The dose makes the poison*” is a founding principle of toxicology ascribed to Swiss physician Paracelsus nearly half a millennium ago, but remains relevant today for understanding both fundamental and applied ways in which organisms interact with chemicals in their environment, including our own medical treatments, dietary regimes, and use of pesticides in agricultural control (Calabrese and Mattson, 2017; Cutler, 2013; Hayes, 2007; Mattson, 2008; Ode and Ghosh, 2025). Our work reveals how an unusual fly species has established a new relationship with an environmental toxin with both individual benefit and potential population impact. How widespread the evolution of hormesis is in animals, whether specialists or generalists, during niche adaptation remains an important future question.

## Acknowledgements

We thank Hector Gallart Ayala and Julijana Ivanisevic at the Metabolomics & Lipidomics Facility (University of Lausanne) for performing the metabolic analyses, Tadeusz Kawecki for advice on PCR-based sexing, and John Pannell and members of the Benton laboratory for discussion and comments on the manuscript. Research in R.B.’s laboratory was supported by the University of Lausanne, an ERC Advanced Grant (833548) and the Swiss National Science Foundation (219185 and 10005666).

## Author contributions

D.M. and R.B. conceived the study. D.M. performed all experiments and analyses with the exception of Figure 4A and Figure S7E (performed by M.M.--B.) and Figure S7F (performed by M.M.). D.M. and R.B. wrote the paper. All authors approved the final version.

## Competing interests

The authors declare no competing interests.

## Data availability

All data and custom codes associated with main and supplementary figures are provided along with the manuscript in Source Data and Source Code.

## METHODS

### *Drosophila* strains and husbandry

All species and strains of drosophilids used in this study (Table S1) were reared in non-overlapping 2-week cycles under 12:12 h light:dark conditions, with 60% humidity at 25°C, unless otherwise specified.

Experiments were performed with 4-7-day old flies, unless otherwise specified. In experiments where the distinction between developmental and adult diet was necessary, adult flies were switched to a different medium within 3 h after eclosion.

### Dietary treatment

A wheat flour-yeast-fruit juice medium (Table S2) served as the “base” medium for all fly rearing. To create control, noni and isocaloric media, this base was supplemented with a ∼3-4 g paste (an excess amount than required to ensure that food availability is not a limiting factor) of instant *Drosophila* medium (Formula 4-24 blue, Carolina Bio-Supply) mixed with tap water, noni juice (Raab Vitalfood Noni Saft) or 20% grape juice (representing an equivalent energy content based upon the manufacturers’ information) (Beutelsbacher Bio Demeter Traubensaft rot), respectively. To test the individual chemical components of noni fruit (Almeida *et al*., 2019; Zhou and Huang, 2024), a single component was directly mixed with base media to the desired final concentration. The base medium density is 1.01778 g/ml.

A nutrient-poor medium for rearing and egg-laying of *D. melanogaster* (Figure 5D-F, H) was prepared by diluting the base medium to 0.25× its original concentration and adding agar (BD Difco^TM^ Agar, 214010) to a final concentration of 0.8% (w/v) to match the consistency and texture of the base medium.

For noni fruit media, ripe noni fruits were collected from *Morinda citrifolia* shrubs in the University of Lausanne greenhouse and stored at −20°C. Whenever fresh noni fruit media was needed, the fruits were thawed, crushed, and the seeds carefully removed. The fruit purée was then placed into empty fly vials, and a small Whatman paper was placed on top of it to prevent flies from sticking to crushed fruit.

### Olfactory behavior

Olfactory behavior of flies was assessed using a planar arena with two odor sources at opposite ends (Maesani et al., 2015). Briefly, a polyoxymethylene and acrylic glass planar arena (50 × 10 × 1.3 mm), illuminated with a far-red-filtered white LED backlight and monitored by a top-mounted acquisition camera (Guppy, Allied Vision Technologies, Stadtroda, Germany), was used to record fly behavior while distinct laminar odor streams were presented at the two ends of the arena. Odor stimulation was generated by bubbling air (Messer Schweiz AG, Lenzburg, Switzerland) through the test odorants (e.g., noni juice, apple cider vinegar, grape juice or water) and delivering the resulting vapor streams at a controlled flow rate of 500 ml/min using mass flow controllers (PKM SA). A custom-fabricated electronic interface and the software sQuid (http://lis.epfl.squid/) were used to control both odor-flow valves and the acquisition camera.

### Feeding rate

The Capillary Feeder (CAFE) assay was used to measure the feeding rate of flies (Diegelmann et al., 2017). Briefly, an empty vial was fitted with a borosilicate glass capillary micropipette (Sutter Instruments, BF100-50-10) filled with either noni juice or a 2.5% sucrose solution (matching the sugar content present in noni juice). The initial liquid level in the capillary was recorded, and ten flies were introduced into the vial. After 2 h, the flies were removed, and the final liquid level was measured with a caliper. The volume of liquid consumed was calculated from the change in liquid level, averaged per fly, and normalized to the average body weight of the flies, measured independently.

### Circadian behavior

Single-fly circadian behavior was assessed using the *Drosophila* Activity Monitor 2 system (DAM2; Trikinetics Inc.) (Luo et al., 2021), following the same setup as for the single-fly starvation resistance assay, except that one end of each fly tube was capped with a 2% agar and 5% sucrose solution to provide continuous access to both water and food. Locomotor activity and sleep parameters were analyzed using custom-written R scripts together with the Rethomics package.

### Male mating success

To assess the effect of diet on male mating success, a 10× mating assay was performed. Briefly, a single 2-4-day-old naïve male was co-housed with ten virgin females for a 24 h period. Each female was then transferred individually to a fresh vial. Starting day 5, the presence of larvae or pupae in a vial was used as an indicator of successful mating. For each successfully mated female, the total number of offspring produced was counted to evaluate the impact of male diet on reproductive output.

### Ovariole numbers

To determine the number of ovarioles, 4-day-old mated females were anesthetized using CO_2_ and transferred to a dissection pad. Under a stereomicroscope, ovaries were carefully dissected in phosphate-buffered saline (PBS) solution using fine forceps. Individual ovarioles were gently separated within each ovary (Wong and Schedl, 2006), and the total number of ovarioles from both ovaries were quantified for each female.

To examine the effect of adult diet on ovariole number, 3-day-old females and males were maintained either on their developmental diet or transferred to an alternative diet (for instance, control-to-control diet or control-to-noni diet). After 1-9 days of dietary exposure, females from each treatment group (control-to-control, control-to-noni, noni-to-noni and noni-to-control) were collected, and their ovariole numbers were determined as described above.

### Egg-laying rates

Egg laying plates were prepared by mixing noni juice, 20% grape juice or water with the base medium (see above) at a 10:90 (v/v) ratio to prepare, noni, isocaloric or control plates, respectively. Ten 4-7-day-old mated females were anesthetized with CO_2_ and transferred to each plate. After 24 h, the females were moved to a fresh plate, and the number of eggs laid on the previous plates was counted manually under a stereomicroscope. This procedure was repeated for three consecutive days to quantify daily egg-laying rates.

For assays involving individual components of noni fruit (Almeida *et al*., 2019; Chan-Blanco et al., 2006; Pino *et al*., 2009; West et al., 2011; Zhou and Huang, 2024), egg-laying plates were prepared by incorporating each component at the required concentration into the control plate medium.

Viability was assessed by counting the number of pupae that developed from eggs laid during the 24 h egg-laying assay. After seven days, all pupae present on each egg-laying plate were counted, and viability was calculated as the ratio of the number of pupae to the total number of eggs laid on that plate.

### Egg and larval fitness

Five mated 4-day-old females were allowed to lay eggs on an egg-laying plate for approximately 24 h. For the egg-transfer assay, eggs were collected and transferred to fresh food vials using a small drop of water placed on a coverslip to facilitate handling. For the larval-transfer assay, plates were incubated for an additional ∼24 h, after which newly hatched larvae were collected and transferred. For both assays, 30 eggs or larvae were transferred into each fresh food vial. After 7 days, the number of pupae in each vial was recorded. After 10 days, emerging adults were counted daily until no additional flies eclosed.

### Reproductive output

*Females*: to measure the reproductive output, five 4-day-old mated females and males (or two of each for generalist species) were transferred to vials containing fresh food medium. After three days, the adults were removed from the vials. Beginning nine days later, emerging offspring were collected daily, sex-sorted, and counted until no additional offspring emerged.

For assays measuring lifetime reproductive output, adult flies were transferred to fresh vials twice per week, and all offspring emerging from each vial were counted over the entire lifespan of the flies.

### Sex ratio analysis

#### Adult

The adult sex ratio was computed by sex-sorting and counting all the adult offspring using the same procedure described above for reproductive output on the media of choice (for instance, control or noni or noni-specific component media). Sex ratio was calculated as the number of female offspring divided by the total number of offspring produced per vial.

For lifetime sex ratio analysis, five females and five males were maintained together for their entire lifespan, transferring to fresh vials every 3 days until all flies had died. Offspring emerging from all vials were sex-sorted and counted, and a cumulative sex ratio was calculated by combining progeny across the entire reproductive period for each replicate.

#### Eggs and larvae

because *Drosophila* eggs/early embryos lack distinguishable morphological sex markers, a PCR-based method was used for sex identification. All eggs laid within a 24-h period in egg laying plate by five 4-day-old mated females were collected individually into the wells of a 96-well plate. DNA extraction was performed by adding 6 μl of squishing buffer (10 mM Tris-HCl, 1 mM EDTA, 25 mM NaCl) supplemented with proteinase K (200 μg/ml) to each well, followed by incubation at 37°C for 30 min and heat inactivation at 95°C for 5 min.

#### Two primer sets were used

1. an autosomal positive-control primer pair targeting *D. sechellia Actin 42A* (LOC6619144; Forward (5’-3’): CAGATGTGGATCTCGAAGCA,

#### Reverse

TTCTGAAGGAGCGAAAGTGT), and (2) a Y-chromosome-specific primer pair targeting *Pp1-Y2* (LOC116803406; Forward: ATCTGCGGTGACATACACGG, Reverse: CGTCACATGGTCGGGCTAAT) to identify male embryos. PCR was performed using KOD One™ PCR Master Mix (Merck) with the following cycling protocol: initial denaturation at 98°C for 2 min; 38 cycles of 98°C for 15 s, 59°C for 10 s, and 68°C for 6 s; final extension at 68°C for 3 min.

To validate the sex-identification assay, both primer sets were initially tested on adult male and female flies, which possess morphologically distinguishable features. We also used similar PCR-based sex-identification assay to confirm sex of larvae.

Larval sex was additionally determined using a genetic cross between males from a *D. sechellia* mutant line carrying an X-linked RFP marker (*w^3P3-RFP^/Y*, generated in a *D. sechellia* 07 background (Auer *et al*., 2020) and wild-type *D. sechellia* 07 females. As a control, we verified that this reporter fly line also exhibited the female-biased sex ratio shift on noni diet in adults (Figure S3H-I).

In the resulting F1 progeny, males inherit a single X chromosome from the wild-type mother and therefore lack fluorescence, whereas females are heterozygous for the RFP insertion and exhibit red fluorescence. Larvae were examined for RFP expression using a Nikon SMZ25 stereomicroscope, and fluorescence was manually scored to assign sex.

For all cases, sex ratio was calculated as the number of female eggs or larvae divided by the total number of eggs or larvae produced per vial.

### Lifespan

Lifespan was determined by keeping five 1-day old females and males together in a vial containing the desired food medium. Ten replicates were used to obtain one set of lifespan data (i.e., 50 female and 50 male flies in total). Live flies were transferred to fresh media vials twice a week. Dead flies and their sexes were recorded every weekday until all the flies had died.

### Starvation and desiccation resistance

Single-fly starvation and desiccation resistance assays were performed using the DAM2 system (see above). Briefly, for starvation assays individual flies were placed in 5-mm glass tubes, with one end capped with 2% agar (providing a water source to prevent desiccation) and the other end sealed with a cotton plug. Thirty-two tubes were loaded into each DAM2 system, and five monitors were housed inside a temperature-controlled incubator maintained at 25°C under a 12:12 h light-dark cycle. The real-time activity of each fly was recorded as the number of infrared beam crossings per minute using DAMsystem311X data acquisition software. The time to death was defined as the last recorded activity event for each individual. All experiments were conducted for six consecutive days. Behavioral data, including locomotor activity, sleep bout analysis, and time of death measurements, were analyzed using custom-written R scripts (available in *Source Code* folder) and the Rethomics package (Geissmann et al., 2019).

Single-fly desiccation resistance assays followed the same recording protocol and analysis as the single-fly starvation resistance assay, except that the tubes did not contain 2% agar. The absence of agar prevented access to water, thereby inducing desiccation stress.

Group starvation resistance assays were performed by anesthetizing 20 4-5-day-old flies with CO_2_ and placing them into vials (25 × 90 mm, Milian SA) containing Kimwipes (Kimtech Science) soaked with 2 ml of water to prevent desiccation. The number of dead flies was recorded every 3 h during daytime hours for a period of four days. For Figure 2C and Figure 4F the data points are reported at 72 h post starvation initiation.

### Stress resistance

Stress resistance was assessed by exposing flies to solutions inducing genotoxic, osmotic, proteotoxic, or oxidative stress (Belyi et al., 2020). Twenty 4-5-day old flies were introduced into a vial (25 × 90 mm, Milian SA) containing Kimwipes (Kimtech Science) saturated with 2 ml of a 5% sucrose solution containing either 15 mM CuSO_4_ (genotoxic stress), 400 mM NaCl (osmotic stress), 5 mM CdCl_2_ (proteotoxic stress) or 18.5 mM paraquat. The number of dead flies was recorded after a 24-h exposure period for the genotoxic, proteotoxic and oxidative stress assays, and after 5 days for osmotic stress assay.

### Heat resistance

Heat resistance was assessed using Critical Thermal Maximum (CT_max_) method (Rokusek et al., 2023). For the CT_max_ assay, 4-5-day-old flies were exposed to gradually increasing temperatures in a water bath. Individual flies were placed in separate vials, which were then submerged in the water bath. A thermometer was placed inside an empty vial under identical conditions to accurately record the temperature experienced by the flies. The temperature at which each fly became immobilized was recorded as the CT_max_.

### Metabolic rate

The metabolic rate of flies was determined using respirometry (Yatsenko et al., 2014). In this assay, CO_2_ production by animals serves as an indirect measure of O_2_ consumption and ATP production via the Krebs cycle; therefore, the amount of CO_2_ produced provides a readout of metabolic rate.

To construct the fly respirometer, a 1000 μl pipette tip was cut and fitted with a borosilicate glass capillary micropipette (Sutter Instruments, BF100-50-10), which was sealed with superglue. A small amount of CO_2_ absorbent (KOH pellet) was placed between two foam pieces inside the pipette tip. Five 4-5-day-old flies were introduced into the chamber, and the open end was sealed with moldable wax. The assembled setup was then placed in a container of dyed water, and the initial water level in the capillary was recorded. As the flies respired, the CO_2_ they produced was absorbed by the CO_2_ absorbent, creating a negative pressure that caused the water to rise in the capillary. After 15 min, the displacement of water was measured to calculate the volume of CO_2_ produced. The volume was averaged per fly and normalized to body weight (measured independently) to estimate the metabolic rate.

### Body weight

For wet weight measurements, groups of 10 flies were anesthetized and transferred into small vials and weighed using a microbalance. For dry weight measurements, groups of 10 flies were placed in microcentrifuge tubes, flash-frozen and dried at 100°C for 24 h before weighing.

### Whole-body bulk metabolomics and analysis

#### Sample preparation

for each biological replicate (*n* = 5) of 10 flies, 4-day-old *D. sechellia* mated adults fed on either the noni or control diet were collected under CO₂ anesthesia, sex-sorted, and flash-frozen in liquid nitrogen.

#### Metabolite extraction protocol

samples were homogenized by the addition of 750 µl of MeOH:H2O (4:1), in the Cryolys Precellys 24 sample Homogenizer (2 × 20 s at 10,000 rpm, Bertin Technologies, Rockville, MD, US) with ceramic beads. The bead beater was air-cooled down at a flow rate of 110 l/min at 6 bar. Homogenized extracts were centrifuged for 15 min at 4000 g at 4°C (Hermle, Gosheim, Germany). Sample extracts were then evaporated to dryness and resuspended in MeOH 80% according to the total protein content (BCA assay).

#### Multiple pathway targeted analysis of polar and moderately polar metabolites

cell lysates were analyzed by hydrophilic interaction liquid chromatography coupled to tandem mass spectrometry (HILIC–MS/MS) using a 6495 triple quadrupole mass spectrometer (QqQ) interfaced with a 1290 UHPLC system (Agilent Technologies, Santa Clara, CA, USA). Two complementary chromatographic modes, coupled to positive and negative electrospray ionization (ESI), were employed to maximize metabolome coverage, encompassing more than 400 intermediates of central carbon metabolism, including glycolysis, the pentose phosphate pathway, purine and pyrimidine metabolism, the tricarboxylic acid (TCA) cycle, amino acid biosynthesis, and oxidative phosphorylation (OXPHOS). Data were acquired in dynamic multiple reaction monitoring (dMRM) mode with a total cycle time of 600 ms, using optimized collision energies for each metabolite. Acquisition was performed with MassHunter Acquisition software (Agilent Technologies) as previously described (van der Velpen et al., 2019).

In positive ionization mode, chromatographic separation was performed on an ACQUITY BEH Amide column (1.7 μm, 100 mm × 2.1 mm i.d.; Waters, Milford, MA, USA). The mobile phase consisted of (A) 20 mM ammonium formate with 0.1% formic acid in water and (B) 0.1% formic acid in acetonitrile. The gradient program was as follows: 0 min, 95% B; 1.5 min, 95% B; 17 min, 45% B; 19 min, 45% B; 20 min, 95% B; and 25 min, 95% B. The flow rate was 400 μl min⁻¹, the column temperature was maintained at 25 °C, and the injection volume was 2 μl. The ESI source parameters were as follows: drying gas temperature, 230°C; drying gas flow, 14 l/min; nebulizer pressure, 35 psi; sheath gas temperature, 400°C; sheath gas flow, 12 l/min; nozzle voltage, 500 V; and capillary voltage, +4000 V.

In negative ionization mode, chromatographic separation was performed on an iHILIC-(P) Classic column (5 μm, 100 mm × 2.1 mm, 200 Å, PEEK; HILICON AB, Umeå, Sweden). The mobile phase consisted of (A) 20 mM ammonium acetate and 20 mM ammonium hydroxide in water (pH 9.45) and (B) 100% acetonitrile. The gradient program was as follows: 0 min, 90% B; 1.5 min, 90% B; 8 min, 50% B; 11 min, 50% B; 12 min, 45% B; 15 min, 45% B; 16 min, 90% B; and 25 min, 90% B. The flow rate was 300 μl/min, the column temperature was maintained at 30°C, and the injection volume was 2 μl. The ESI source parameters were as follows: drying gas temperature, 290°C; drying gas flow, 14 l/min; nebulizer pressure, 45 psi; sheath gas temperature, 350°C; sheath gas flow, 12 l/min; nozzle voltage, 0 V; and capillary voltage, −2000 V.

For relative quantification of polar metabolites, raw LC-MS/MS data were processed using MassHunter Acquisition software (Agilent Technologies, Santa Clara, CA, USA). Relative quantification of metabolites was based on extracted ion chromatogram (EIC) areas corresponding to the monitored MRM transitions. Signal intensity drift over time was corrected for each annotated metabolite feature by fitting a locally weighted scatterplot smoothing (LOWESS) curve to quality control (QC) pooled samples, which were representative of the entire dataset and analyzed periodically throughout the batch. Data were further curated by removing peaks with high analytical variability, defined as a coefficient of variation across QC samples greater than 30%.

#### Untargeted metabolomics analysis of polar metabolites

metabolite relative quantification was performed using ultra-high performance liquid chromatography coupled to high resolution mass spectrometry instrument (UHPLC-HRMS). Data acquisition was performed in full scan mode on Tribrid IQ-X Orbitrap (Thermo Fisher Scientific) instrument operating at mass resolving power of 120K. Two complementary chromatographic separations coupled to positive and negative electrospray ionization MS were used to maximize metabolome coverage. For metabolite annotation AcquireX workflow in ddMS^2^ mode was applied. Metabolite fragmentation was obtained at three collision energies (stepped 10, 30 and 55 NCE).

In positive mode, the chromatographic separation was carried out in an Acquity BEH Amide, 1.7 μm, 100 mm × 2.1 mm i.d. column (Waters, Massachusetts, US). Mobile phase was composed of A = 20 mM ammonium formate and 0.1 % FA in water and B = 0.1 % FA in ACN. The linear gradient elution from 95% B (0-1.5 min) down to 45% B was applied (1.5 min −17 min) and these conditions were held for 2 min. Then initial chromatographic conditions were maintained as a post-run during 5 min for column re-equilibration. The flow rate was 400 μl/min, column temperature 25°C and sample injection volume 2 µl.

In negative mode, chromatographic separation was performed on an iHILIC-(P) Classic column (5 μm, 100 mm × 2.1 mm, 200 Å, PEEK; HILICON AB, Umeå, Sweden). The mobile phase consisted of (A) 20 mM ammonium acetate and 20 mM ammonium hydroxide in water (pH 9.45) and (B) 100% acetonitrile. The gradient program was as follows: 0 min, 90% B; 1.5 min, 90% B; 8 min, 50% B; 11 min, 50% B; 12 min, 45% B; 15 min, 45% B; 16 min, 90% B; and 25 min, 90% B. The flow rate was 300 μl/min, the column temperature was maintained at 30 °C, and the injection volume was 2 μl.

For data processing, raw LC-HRMS(/MS) data were processed using Compound Discoverer^TM^ 3.3 software (Thermo Fisher Scientific). Relative quantification of metabolites was based on EIC (Extracted Ion Chromatogram) areas for the m/z. The obtained tables (containing peak areas of detected metabolites across all samples) were processed and filtered according to the variability of each metabolite in pooled QC samples (representatives of the entire data set). Pooled QC samples were analyzed periodically throughout the overall analytical run and used to assess the quality of the data and correct the signal intensity drift (attenuation in most cases, that is inherent to LC-MS technique and MS detector due to sample interaction with the instrument over time) and remove the peaks with poor reproducibility (CV > 30%). Metabolite annotation was performed using Thermo Scientific mzCloud spectral library and the identification of the compounds was made according to the following criteria: retention time alignment, mass error <5 ppm for precursor ions with a minimum of one fragment ion, the MS^2^ spectra comparison between the observed theoretical fragmentation followed by the mzCloud database having a MS/MS matching score higher than 65% and mass error<5ppm.

#### Protein quantification

The protein pellets were evaporated and lysed in 20 mM Tris-HCl (pH 7.5), 4 M guanidine hydrochloride, 150 mM NaCl, 1 mM Na_2_EDTA, 1 mM EGTA, 1% Triton, 2.5 mM sodium pyrophosphate, 1 mM beta-glycerophosphate, 1 mM Na_3_VO_4_, 1 µg/ml leupeptin using the Cryolys Precellys 24 sample homogenizer (2 × 20 s at 10,000 rpm, Bertin Technologies, Rockville, MD, US) with ceramic beads. BCA Protein Assay Kit (Thermo Scientific, Massachusetts, US) was used to measure (A562 nm) total protein concentration (Hidex, Turku, Finland).

The statistical, pathway and enrichment analysis were performed using Metaboanalyst 6.0 (www.metaboanalyst.ca) on the log-transformed pareto-scaled metabolite abundance data for both the MPTA and the untargeted metabolomics.

### Macronutrient analysis

For protein and glucose quantification, ten 4-5-day old flies were anesthetized with CO_2_, sex-sorted, weighed, and flash-frozen. The flies were homogenized using a motorized pestle mixer in 1 ml of 0.05% Tween-20 in PBS (Li et al., 2016). The homogenates were centrifuged at 4000 rpm for 30 s to remove debris, and the supernatants were collected in fresh tubes. Total protein content was quantified using the Pierce^TM^ BCA Protein Assay Kit (Thermo Scientific), and total glucose content was measured using the Glucose (GO) Assay Kit (Sigma-Aldrich) (Schwasinger-Schmidt et al., 2012). Each sample was assayed in triplicate, and values were averaged across three technical replicates.

For lipid quantification, ten 4-5-day old flies were anesthetized with CO_2_, sex-sorted, weighed, and flash-frozen. The flies were dried at 100°C for 24 h to obtain dry weight. Lipids were extracted by adding diethyl ether to each tube, followed by gentle agitation on a rocker for 24 h. After ether evaporation, the samples were again dried at 100°C for 24 h and weighed. Total lipid content, expressed as a fraction of dry weight, was calculated as: 1 - (post-extraction weight/dry weight) (Ballard *et al*., 2008).

### Fat body staining and lipid droplet quantification

#### Whole-mount staining and confocal imaging

4-day-old adult females were collected, and fat body were dissected from the abdominal carcasses in 1× PBS after removal of the gut and reproductive organs. The dissected carcasses were pinned onto Sylgard-coated wells, with the internal fat body exposed, and fixed in 4% paraformaldehyde in PBS for 30 min at room temperature. Samples were washed twice for 15 min in PBS containing 0.1% Triton X-100 (PBST). F-actin was stained with Phalloidin-iFluor 594 (Abcam ab176757) for 2 h at room temperature, protected from light, followed by a 15-min PBST wash. Lipid droplets were subsequently stained with BODIPY 493/503 (Thermo Fisher Scientific, D3922) at 1-2 µg/ml in PBS for 15-30 min at room temperature in the dark. Following a final 15-min wash in PBS, samples were mounted in DAPI-containing mounting medium (VECTASHIELD Antifade Mounting Medium with DAPI, H-1200-10) and stored at 4°C, protected from light, until imaging (Ott et al., 2024). Confocal images were acquired at 63× magnification with oil immersion using Zeiss LSM 880 Airyscan with identical acquisition settings (Zen black 2.1 SP3) across all experimental conditions.

#### Image acquisition and quantification

Images were analyzed using custom python scripts. Three parameters were quantified from the BODIPY channel: total fluorescence intensity, lipid-droplet number and lipid-droplet size. Background-corrected integrated BODIPY fluorescence intensity was measured within the analyzed fat-body region of interest (ROI). Background signal was estimated from an adjacent region devoid of tissue and subtracted from the integrated signal. The analyzed tissue area was kept consistent across images or accounted for during normalization. Lipid droplets were identified by segmentation of the BODIPY signal using a consistent image-analysis procedure across samples. Droplet number was quantified within the analyzed ROI and normalized to tissue ROI size. Droplet size was quantified as the area of individual segmented droplets, expressed in µm². Droplet-level measurements were summarized for each biological replicate before statistical analysis, with the biological replicate, rather than each individual droplet, treated as the experimental unit. Imaging and analysis parameters were kept consistent across experimental conditions.

### Population dynamics model and sensitivity analysis

#### Model structure

We developed an exploratory, discrete-time, stochastic two-sex population model to investigate how fecundity, offspring sex ratio and survival-related effects can jointly influence population dynamics. Simulations were implemented in Python using NumPy. Male (Mₜ) and female (Fₜ) abundances were tracked separately at each integer time step t, with total population size defined as Nₜ = Mₜ + Fₜ. Simulations ran for up to T = 200 time steps and terminated earlier if the population became extinct.

The baseline model assumed an initial population of N₀ = 200 adults, split equally between males and females, and a density-dependence scale parameter K = 1,000. Baseline birth and mortality parameters were b = 0.30 and d = 0.125, respectively. Male fertilization capacity was set to c = 1.5 females per male per time step, and the strength of density dependence was set to α = 1.0. These parameter values and the treatment-effect multipliers were illustrative rather than fitted directly to experimentally measured vital rates; the simulations therefore explore potential demographic consequences rather than provide quantitative forecasts of natural populations.

#### Environmental stochasticity

Environmental variation in birth and mortality rates was modelled using separate, temporally autocorrelated stochastic processes. For each process, environmental deviations followed an autoregressive process of order one:

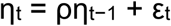

The innovations were normally distributed, with variance determined by the process-specific standard deviation σ and autocorrelation coefficient ρ:

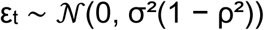

The initial environmental deviation was drawn from a normal distribution with mean zero and standard deviation σ. The baseline standard deviations were 0.20 for births (σ_b_) and 0.10 for mortality (σ_d_), with temporal autocorrelation ρ = 0.65 for both processes.

Birth rates were perturbed multiplicatively using lognormal environmental variation. The fecundity multiplier f_b_ scales the baseline birth parameter:

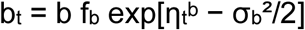

Mortality probabilities were perturbed on the logit scale. The multiplier fᵈ is applied to the baseline mortality parameter before clipping it to the interval [10⁻⁶, 1 − 10⁻⁶]:

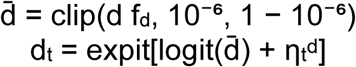

Here, logit(x) = ln[x/(1 − x)] and expit(x) = 1/[1 + exp(−x)]. In the code, this multiplier is labelled “survival”; values below one reduce the baseline mortality parameter and values above one increase it. The same realized mortality probability was applied to males and females; the model does not include a sex-specific survival effect.

#### Reproduction, density dependence and population updates

Reproduction was constrained by male availability. When both sexes were present, the fraction of females fertilized was calculated from male fertilization capacity c and the current sex-specific abundances:

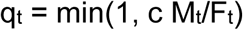

If either sex was absent, the fertilized fraction was set to zero. Expected births were then calculated from the effective birth rate, the number of females, the fertilized fraction and a density-dependent recruitment multiplier:

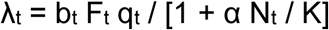

Realized total births were drawn from a Poisson distribution with mean λₜ. The fraction of female offspring s determined sex assignment:

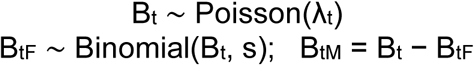

Male and female deaths were sampled independently from binomial distributions using their respective abundances and the shared mortality probability dₜ:

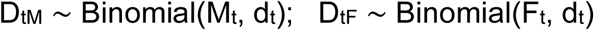

Population abundances were updated by adding births and subtracting deaths:

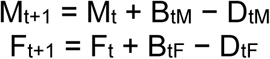

Density dependence acts on recruitment through the multiplier 1/[1 + αNₜ/K]; population size is not rescaled to impose a hard carrying-capacity limit.

#### Simulation design and demographic analyses

To examine the individual effects of offspring sex ratio and fecundity, we simulated population trajectories across offspring female fractions from 0.40 to 0.70 in increments of 0.05, and fecundity multipliers from 0.25 to 1.75 in increments of 0.25. Each condition was evaluated using 100 stochastic replicates. For the joint analysis, we constructed a 100 × 100 parameter grid spanning fecundity multipliers of 0.1-2.0 and offspring female fractions of 0.35-0.85. Each grid point was evaluated using 30 stochastic simulations.

Population growth was quantified as mean log population growth per time step. The starting and final population sizes were floored at one to avoid undefined logarithms when a population went extinct, and the denominator was the elapsed simulation duration:

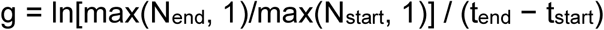

The heatmap displays the mean growth metric across replicates at each parameter combination. Trajectory plots display mean population size with the interquartile range (25th–75th percentiles) across replicates.

We used Shapley decomposition to quantify the contributions of fecundity, offspring sex ratio and the mortality multiplier to changes in population growth. The control scenario used a fecundity multiplier of 1.0, an offspring female fraction of 0.50 and a mortality multiplier of 1.0. The illustrative combined-treatment scenario used a fecundity multiplier of 1.25, an offspring female fraction of 0.60 and a mortality multiplier of 0.95. For each of the eight possible combinations of the three components, mean log population growth was estimated from 200 stochastic simulations. The marginal contribution of each component was calculated across all possible orderings of the three components and averaged to obtain its Shapley contribution. The same set of replicate seeds was used across component combinations to reduce Monte Carlo noise. Population trajectories for the control, fecundity-only, sex-ratio-only, mortality-only and combined scenarios were evaluated separately using 60 replicates per condition.

#### One-at-a-time sensitivity analysis

We assessed the sensitivity of mean log population growth to 14 model parameters. Each parameter was evaluated at five values while all other parameters were held at their baseline settings. Forty stochastic replicates were simulated for each parameter value. For each parameter setting, we calculated mean log population growth per time step, its standard deviation across replicates, mean final population size and the proportion of replicates ending in extinction. Sensitivity plots display mean log population growth with shaded bands indicating ±1 standard deviation. All simulations used a fixed base random seed of 42, with deterministic seed offsets across conditions and replicates for reproducibility.

### Statistical analysis

All statistical analyses were performed in R/RStudio (4.5.0) and GraphPad Prism 10. The specific statistical tests, models, and associated p-values used for each experiment are reported in the corresponding figure legends. The figure legends also indicate the number of biological replicates and, where applicable, technical replicates used in each assay. No data points were excluded unless explicitly stated.

## Code availability

All MATLAB, Python and R scripts used to perform all simulations, data and statistical analysis and generate figures are provided in the Supplementary Information.

## AI-use statements

Text language refinement and code editing were assisted using ChatGPT (OpenAI; GPT-5.2).

## Supplementary Material

### Supplementary Tables

**Table S1.** *Drosophila* strains.

| **Genotype** | **Reference/Source** |
|---|---|
| *D. sechellia* 07 (*Dsec*07) | *Drosophila* Species Stock Center (DSSC):14021-0248.07 |
| *D. sechellia* 28 (*Dsec*28) | DSSC:14021-0248.28 |
| *D. sechellia w<sup>3P3-RFP</sup>*/*w<sup>3P3-RFP</sup>* | (Auer et al., 2020) |
| *D. sechellia Alkbh7<sup>RFP</sup>* | (Marconcini *et al*., 2026a) |
| *D. simulans* 04 (*Dsim*04) | DSSC:14021-0251.004 |
| *D. simulans* 196 (*Dsim*196) | DSSC:14021-0251.196 |
| *D. melanogaster* Canton-S (*Dmel*CS) |  |
| *D. melanogaster* Oregon R (*Dmel*OR) |  |
| *D. melanogaster Alkbh7<sup>1</sup>* | (Marconcini *et al*., 2026a) |

**Table S2.** Base medium composition.

| **Component** | **Amount** | **Source** |
|---|---|---|
| Agar | 6.2 g | Brunschwig #400400050 |
| Farigel wheat | 58.8 g | Westhove #FMZH1 |
| Yeast | 58.8 g | Springaline BA10 Springer #510 |
| Fruit juice (multivitamin) | 100 ml | Coop |
| Propionic acid | 4.9 ml | Sigma-Aldrich P1386 |
| Moldex (400 g methyl-4-hydroxybenzoate in 4 l 95% ethanol) | 26.5 ml | VWR #ALFAA14289.0 |
| Water | make up to 1 l |  |

**Table S3.** Chemical list.

| Chemical | Reference/source | CAS |
|---|---|---|
| Acetic acid | FLUKA 45726 | 64-19-7 |
| Butyric acid | SIGMA-ALDRICH B103500 | 107-92-6 |
| Isovaleric acid | FLUKA 59850 | 503-74-2 |
| Hexanoic acid | SIGMA-ALDRICH 153745 | 142-62-1 |
| Heptanoic acid | FLUKA 75190 | 111-14-8 |
| Octanoic acid | SIGMA-ALDRICH C2875 | 124-07-2 |
| Nonanoic acid | FLUKA 76342 | 112-05-0 |
| 2-ethylhexanoic acid | ALDRICH 538701 | 149-57-5 |
| *1*-octanol | FLUKA 74848 | 111-87-5 |
| *2*-octanone | SIGMA-ALDRICH O4709 | 111-13-7 |
| Ethyl hexanoate | SIGMA-ALDRICH 148962 | 123-66-0 |
| Octanal | SIGMA-ALDRICH O5608 | 124-13-0 |
| Sucrose | SIGMA-ALDRICH S5690 | 57-50-1 |
| D(+)-Glucose anhydrous | NeoFroxx 1179 | 50-99-7 |
| D(−)-Fructose | SIGMA-ALDRICH F0127 | 57-48-7 |
| L-Dopa-(phenyl-d3) | ALDRICH 333786 | 53587-29-4 |
| Scopoletin | SIGMA-ALDRICH S2500 | 92-61-5 |
| Tween 20 | APPLICHEM A4974.0500 |  |
| Ethanol | CARLO ERBA 4146082 | 64-17-5 |
| Diethyl ether | CARLO ERBA REAGENTS 447521 | 60-29-7 |
| Potassium hydroxide | SIGMA-ALDRICH P1767 | 1310-58-3 |
| Cadmium chloride | SIGMA-ALDRICH 439800 | 10108-64-2 |
| Sodium chloride | SIGMA-ALDRICH 204439 | 7647-14-5 |
| Copper(II) sulfate pentahydrate | SIGMA-ALDRICH C7631 | 7758-99-8 |
| phalloidin-iFluor 594 | ABCAM ab176757 |  |
| BODIPY 493/503 | THERMO FISHER SCIENTIFIC D3922 |  |
| VECTASHIELD Antifade Mounting Medium with DAPI | VECTOR LAB H-1200-10 |  |

### Supplementary Figure Legends

**Figure S1.**
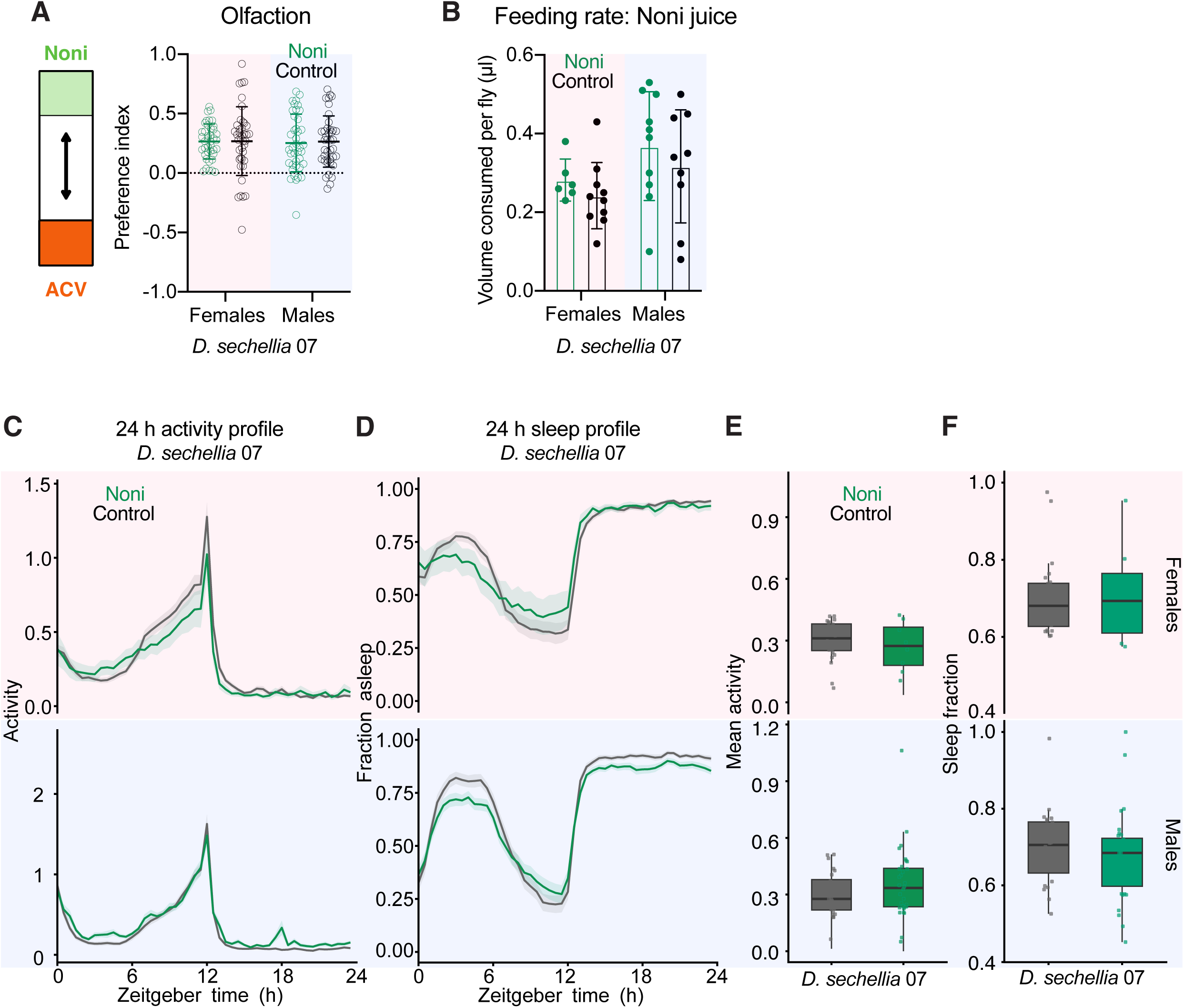
Influence of a noni diet on behavior of specialist *D. sechellia*. (A) Preference index for noni juice versus apple cider vinegar for a two-choice olfactory assay using the planar arena for *D. sechellia* 07 reared on noni or control diet (*n* ≥ 40). (B) Feeding rate computed by measuring the volume of undiluted noni juice (100%) solution consumed per fly using the CAFE assay (*n* ≥ 5 biological replicates of 10 fed flies each). (C-F) Locomotor and circadian behavior for females (top) and males (bottom) *D. sechellia 07* fed on noni or control diet: (C) 24-h locomotor activity profile, (D) 24-h sleep profile, (E) overall mean activity and (F) sleep fraction (*n* ≥ 8 flies). Data in (A–B) and (E–F) were analyzed using two-way ANOVA, with diet and sex as factors, followed by Šídák’s multiple-comparisons test to compare diets within each sex. Significance levels are indicated as \*\*\*\**P* < 0.0001, \*\*\**P* < 0.001, \*\**P* < 0.01, \**P* < 0.05; non-significant comparisons are not labeled.

**Figure S2.**
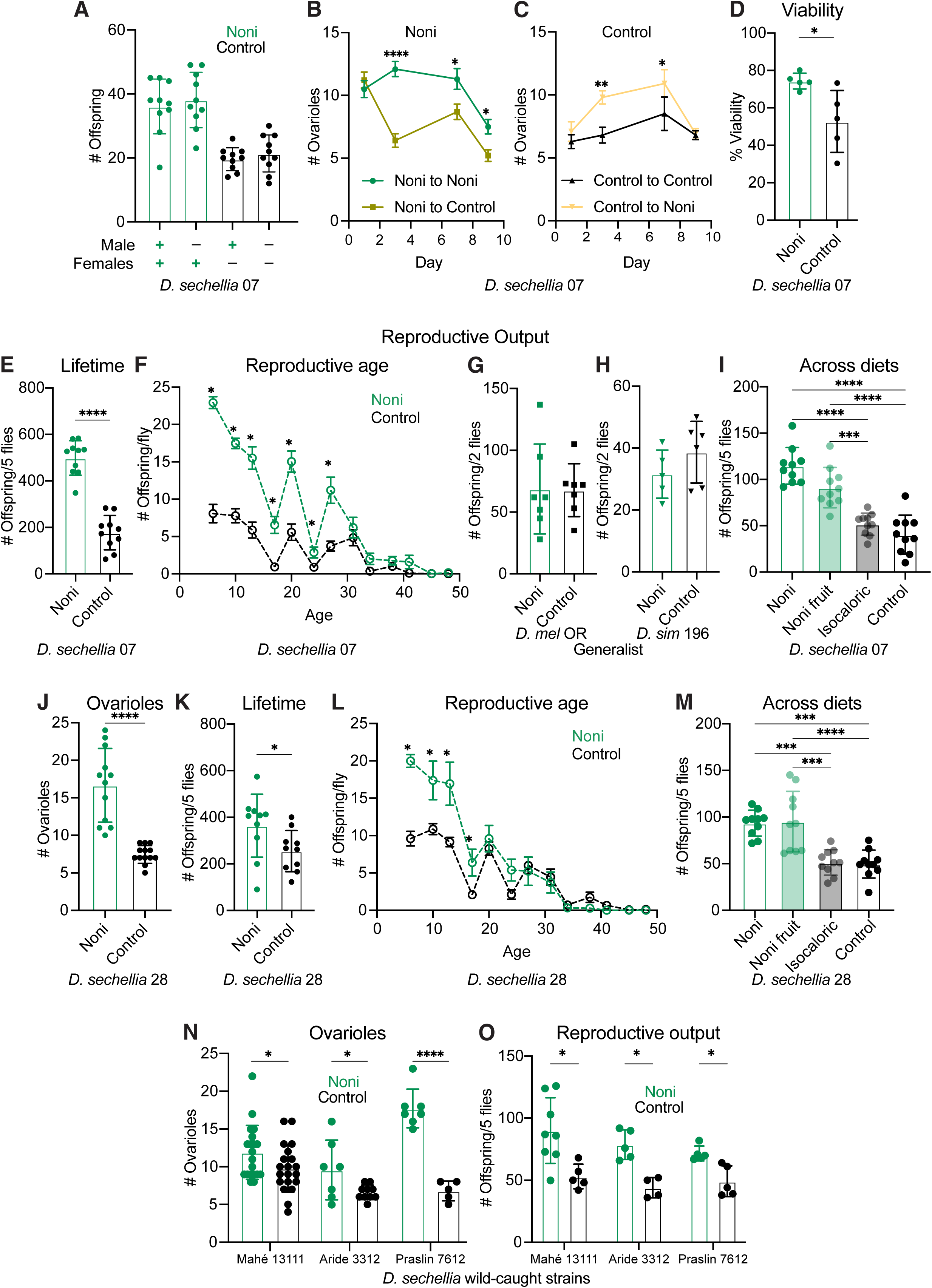
Influence of noni diet on reproductive and developmental traits. (A) Effects of male and female diets (noni or control diet) on reproductive output, quantified as the number of offspring produced (*n* ≥ 10). (B-C) Mature ovariole number over 9 days following diet switches (C) noni to control (or maintaining on noni), and (D) control to noni (or maintaining on control) (*n* ≥ 10 females). (D) Viability for *D. sechellia* 07 embryos on noni or control diet (*n* ≥ 5). (E) Total lifetime reproductive output for *D. sechellia* 07 reared on noni or control diet (*n* ≥ 10 replicates with 5 females each). (F) Offspring per female fly over the lifetime for *D. sechellia* 07 reared on noni or control diet (*n* ≥ 10 replicates with 5 females each). (G-H) Reproductive output for generalist (G) *D. melanogaster* Oregon-R and (H) *D. simulans* 196 reared on noni or control diet (*n* ≥ 5). Two females were used per replicate (instead of five females used in *D. sechellia* experiments) due to higher fecundity of these species. (I) Reproductive output across diets (noni diet, crushed noni fruit, isocaloric grape juice supplemented diet and control diet) for *D. sechellia* 07 (*n* ≥ 10 replicates with 5 females each). (J-M) Reproductive fitness of another independent laboratory strain of *D. sechellia* reared on a noni or control diet. (J) Mature ovariole number (*n* ≥ 10 females), (K) total lifetime reproductive output (*n* ≥ 10 replicates with 5 females each), (L) offspring per female fly over the lifetime (*n* ≥ 10 replicates with 5 females each), and (M) reproductive output across diets (noni diet, crushed noni fruit, isocaloric grape juice supplemented diet and control diet) (*n* ≥ 10 replicates with 5 females each). (N-O) Reproductive fitness of wild-caught *D. sechellia* strains from the Seychelles reared on noni or control diet. (N) Mature ovariole number (*n* ≥ 7 females per strain) and (O) reproductive output (*n* ≥ 5 replicates with 5 females each). Data in (A) were analyzed using a generalized linear model (GLM) with male diet, female diet and their interaction as factors, assuming a Poisson error distribution. Data in (B-C) were analyzed using two-way ANOVA, with diet treatment and day as factors, followed by Šídák’s multiple-comparisons test to compare diet treatments at each day. Data in (D), (E), (G-H), (J-K), and (N-O) were analyzed using two-sided Mann–Whitney U tests to compare control and noni diets within each strain. Data in (F) and (L) were analyzed using a negative-binomial generalized linear mixed model (GLMM), with diet, age and their interaction as fixed effects and replicate identity as a random effect to account for repeated measurements. Šídák-adjusted comparisons were used to compare diets at each age. Data in (I) and (M) were analyzed separately using one-way ANOVA, followed by Tukey’s multiple-comparisons test for all pairwise comparisons between diets. Significance levels are indicated as \*\*\*\**P* < 0.0001, \*\*\**P* < 0.001, \*\**P* < 0.01, \**P* < 0.05; non-significant comparisons are not labeled.

**Figure S3.**
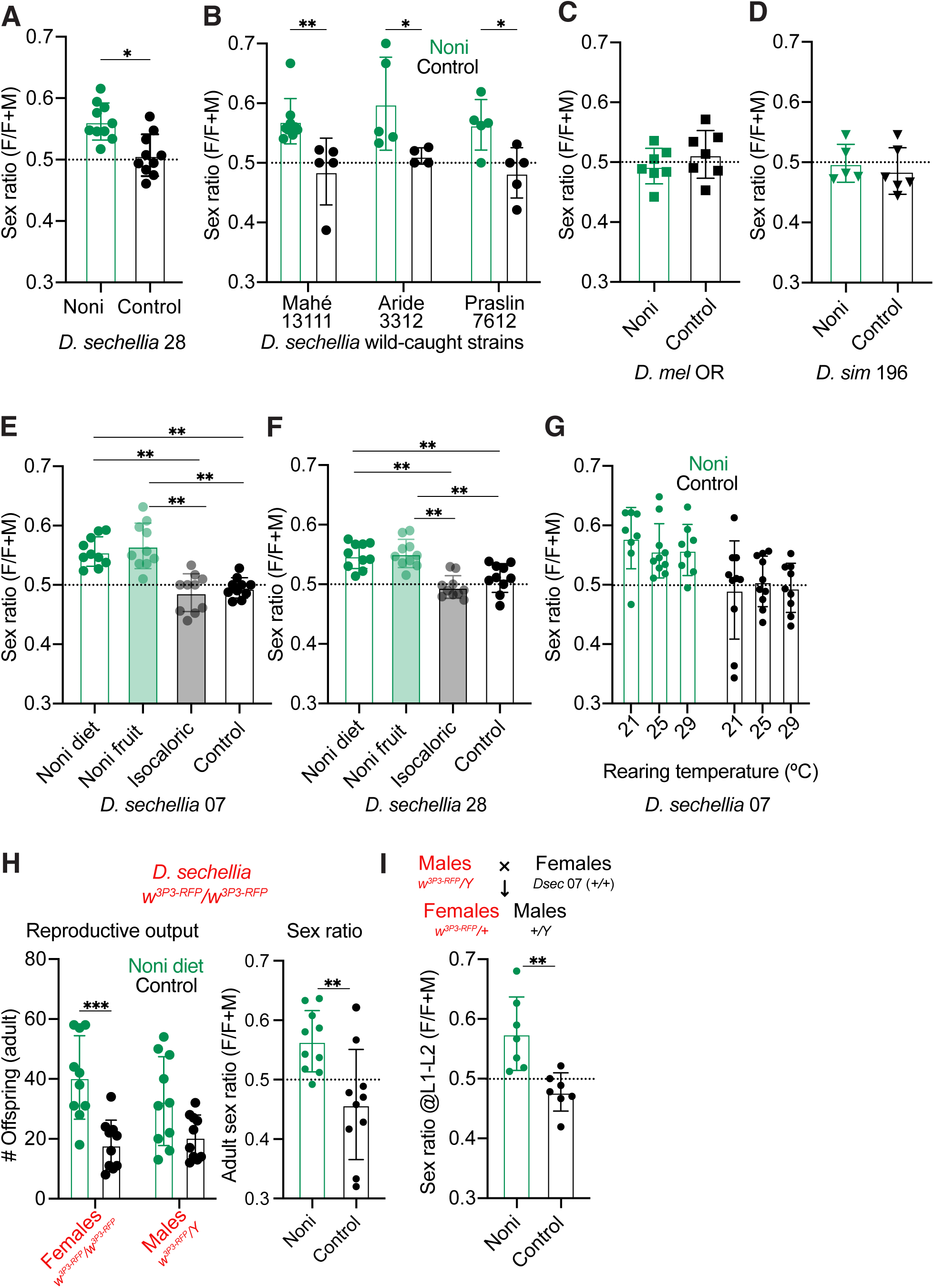
Influence of noni on sex ratio. (A-B) Offspring sex ratio (female offspring/total offspring) for (A) laboratory strain *D. sechellia* 28 (*n* = 10 females) (B) wild-caught strains from the Seychelles (*n* ≥ 7 replicates per strain) reared on a noni or control diet. (C-D) Offspring sex ratio for generalist *D. melanogaster* Oregon-R and *D. simulans* 196 reared on a noni or control diet (*n* ≥ 5 replicates). (E-F) Offspring sex ratio across diets (noni diet, crushed noni fruit, isocaloric grape juice supplemented diet and control diet) for (E) *D. sechellia* 07 and (F) *D. sechellia* 28 (*n* ≥ 10 replicates with 5 females each). (G) Offspring sex ratio at different rearing temperature for *D. sechellia* 07 reared on a noni or control diet (*n* ≥ 5 replicates). (H) Reproductive output (left) and offspring adult sex ratio (right) for *D. sechellia* mutant line carrying an X-linked RFP marker (*D. sechellia* 07 *white^3P3-RFP^/white^3P3-RFP^*) reared on a noni or control diet (*n* ≥ 10 replicates). (I) Larval offspring sex ratio of *D. sechellia* 07 reared on control or noni diet, determined using a genetic cross between males carrying an X-linked RFP marker and wild-type females. In the resulting F1 progeny, females inherit the RFP-marked paternal X chromosome and fluoresce red, whereas males inherit the paternal Y chromosome and the unmarked maternal X chromosome and do not fluoresce (*n* ≥ 7 replicates). Data in (A-D), (H) and (I) were analyzed using two-sided Mann–Whitney U tests to compare control and noni diets within each strain or species. Data in (E-F) were analyzed using one-way ANOVA, followed by Tukey’s multiple-comparisons test for all pairwise comparisons among diets. Data in (G) were analyzed using two-way ANOVA, with diet and temperature as factors, followed by Šídák’s multiple-comparisons test to compare temperature for each diet. Significance levels are indicated as \*\*\*\**P* < 0.0001, \*\*\**P* < 0.001, \*\**P* < 0.01, \**P* < 0.05; non-significant comparisons are not labeled.

**Figure S4.**
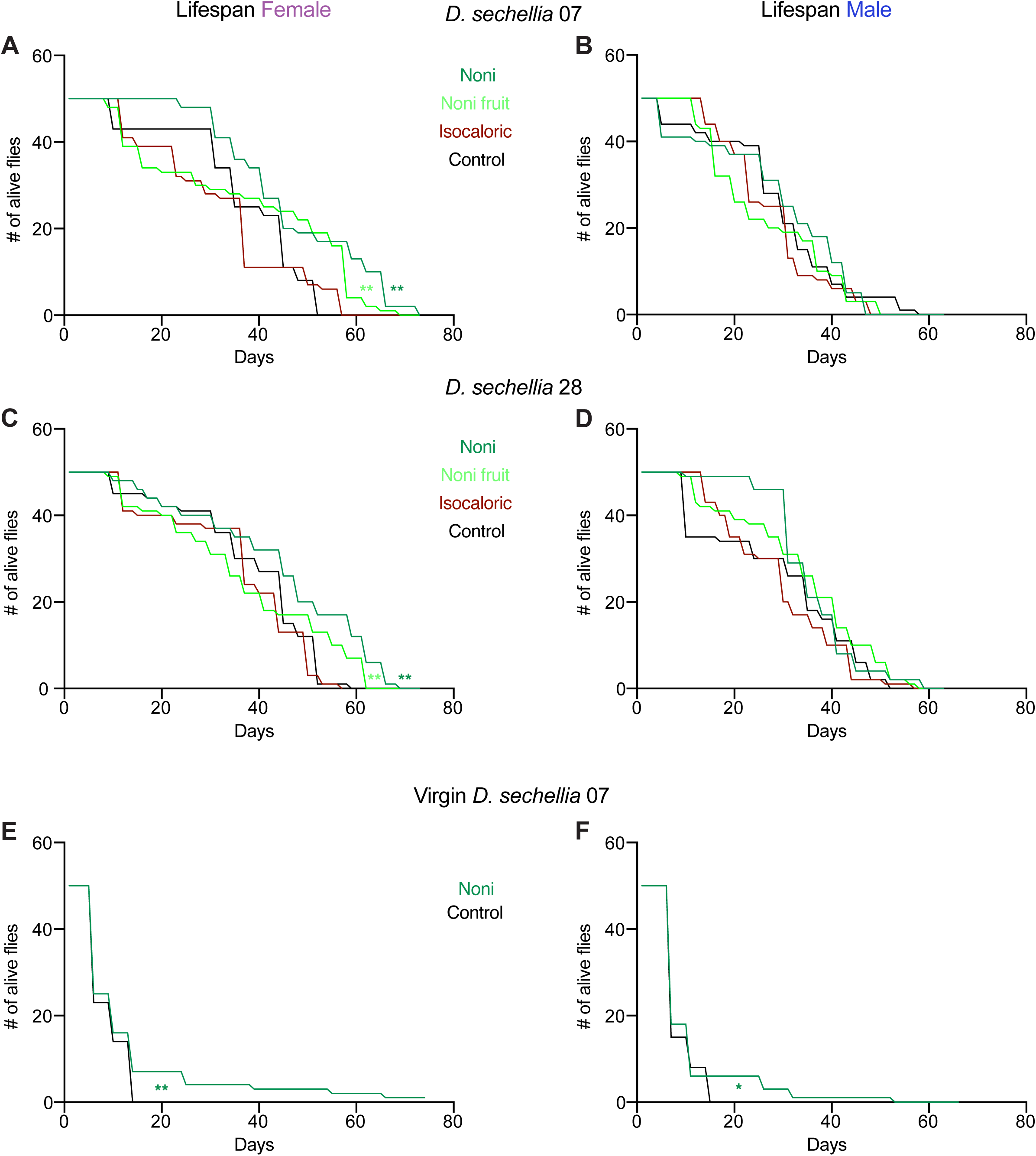
Influence of noni on lifespan. (A,B) Lifespan of (A) females and (B) males of *D. sechellia* 07 maintained on noni juice, crushed noni fruit, isocaloric grape juice supplemented, or control diets. (C,D) Same as (A,B) for *D. sechellia* 28. (E,F) Lifespan of (E) females and (F) males of virgin *D. sechellia* 07. For all panels, *n* = 50 flies per condition. Survival curves were compared using log-rank and Cox proportional hazards tests. Significance levels are indicated as **** *P* < 0.0001, *** *P* < 0.001, ** *P* < 0.01, * *P* < 0.05; non-significant comparisons are not labeled.

**Figure S5.**
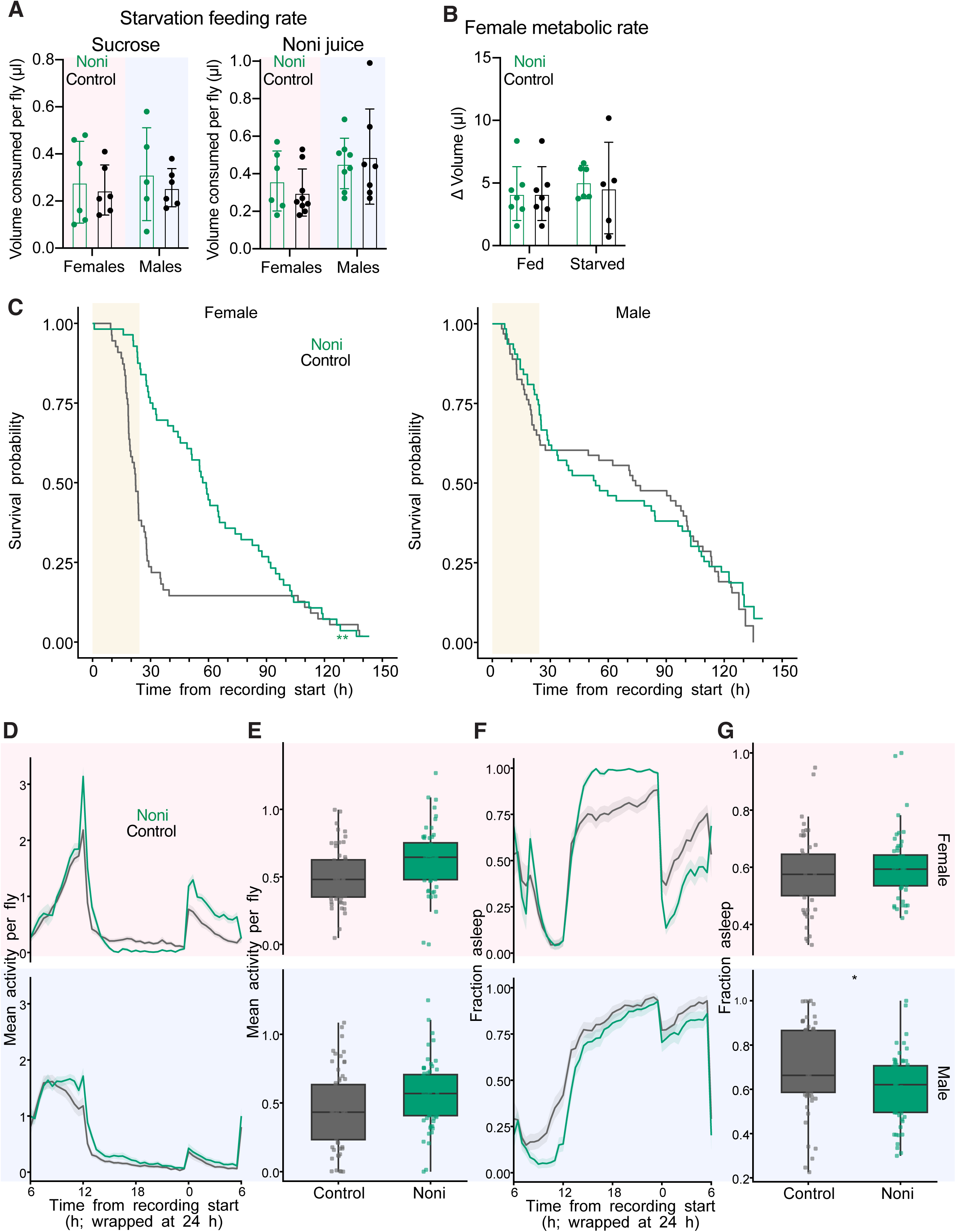
Influence of noni on heat and starvation resistance. (A) Feeding rate for 24-h starved flies for sucrose (left) and undiluted noni juice (100%) (right) consumed per fly using the CAFE assay (*n* ≥ 5 biological replicates of 10 flies each). Note the feeding rate of fed flies under same conditions are already presented in Figure 1C and Figure S1B (*n* ≥ 5). (B) Metabolic rate of fed and starved female *D. sechellia* 07 reared on a noni or control diet (*n* ≥ 6). (C) Survival curves under starvation for *D. sechellia* 07 females (left) and males (right) reared on a noni or control diet (*n* ≥ 55). Shaded regions represent the initial 24 h period used to analyze activity and sleep behavior in (E-H). (D-G) Locomotor and circadian behavior for females (top) and males (bottom) *D. sechellia 07* fed on noni or control diet: (E) 24-h locomotor activity profile, (F) overall mean activity, (G) 24-h sleep profile and (H) sleep fraction (*n* ≥ 55 flies). Data in (A), (E) and (G) were analyzed using two-way ANOVA, with diet and sex as factors, followed by Šídák’s multiple-comparisons test to compare diets within each sex. Data in (B) were analyzed using two-way ANOVA, with diet and starvation state as factors, followed by Šídák’s multiple-comparisons test to compare diets within each starvation state. Survival curves in (C) were compared using log-rank and Cox proportional hazards tests. Significance levels are indicated as **** *P* < 0.0001, *** *P* < 0.001, ** *P* < 0.01, * *P* < 0.05; non-significant comparisons are not labeled.

**Figure S6.**
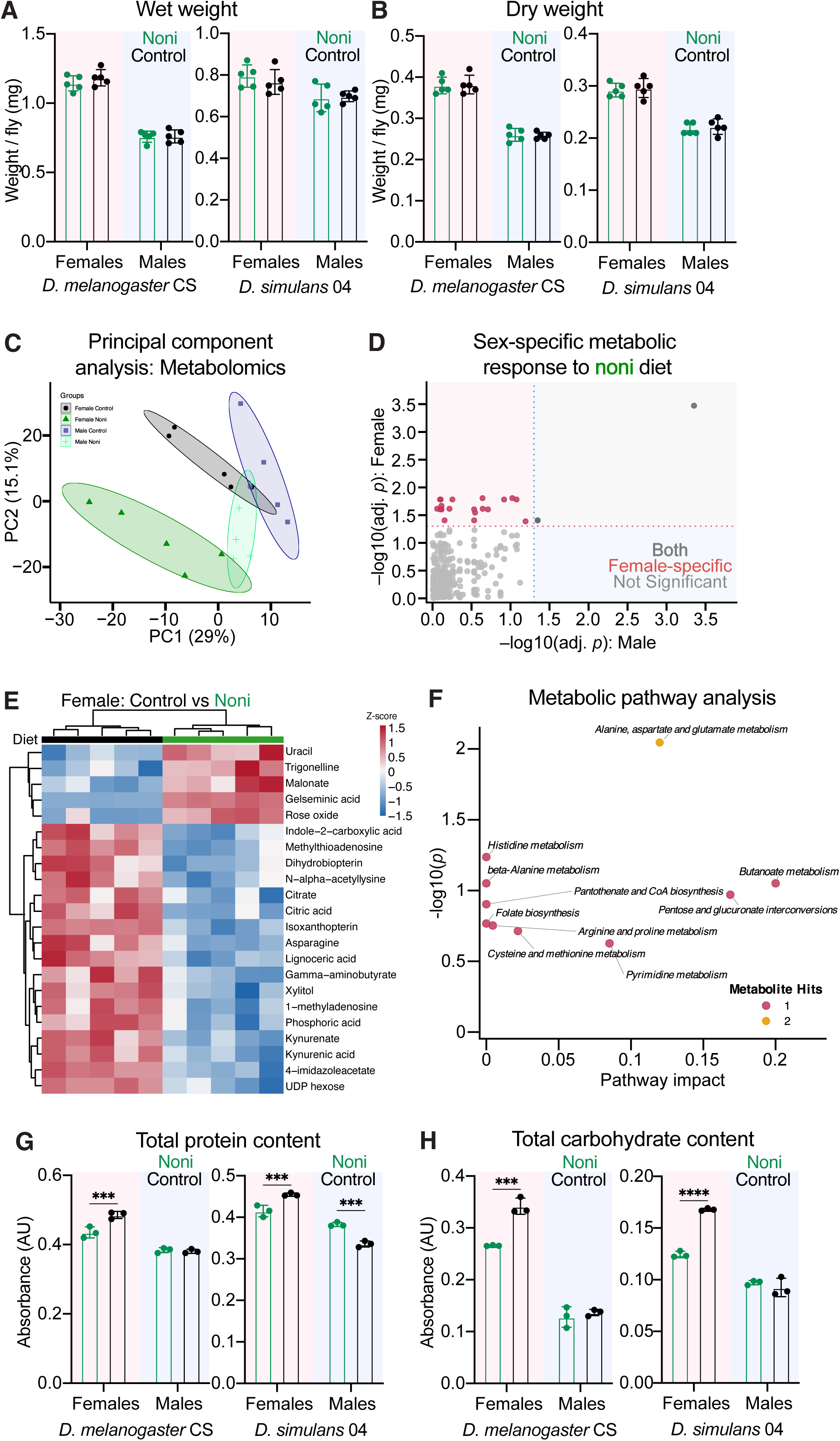
Influence of noni on the *D. sechellia* metabolome and generalist macromolecule composition. (A) Wet body weight of *D. melanogaster* CS and *D. simulans* 04 females and males on noni or control diet (*n* = 5 biological replicates of 10 flies each). (B) Dry body weight of *D. melanogaster* CS and *D. simulans* 04 females and males on noni or control diet (*n* = 5 biological replicates of 10 flies each). (C) Principal component analysis of the whole-body metabolomics (multi-pathway targeted and untargeted analysis combined) dataset for *D. sechellia* 07 females and males on noni or control diet. *n* = 5 samples per sex per diet treatment. (D) Dual volcano plot of noni diet-induced metabolite changes: −log₁₀(adj. *P*) females (y-axis) vs. males (x-axis). Colors: female-specific (pink), male-specific (blue), both (grey), neither (light grey). Dotted lines denote FDR 0.05 thresholds on each axis. (E) Heatmap of relative abundance 22 differentially abundant metabolites in female *D. sechellia* 07 following noni feeding (*n* = 5 biological replicates per diet). Rows represent metabolites and columns represent individual biological replicates. Abundance values were z-score transformed separately for each metabolite across all samples. (F) Metabolic pathway analysis (MetaboAnalyst 6.0) on the noni diet-induced female-specific metabolite changes (pink dots in (D)). (G) Total protein content of *D. melanogaster* CS and *D. simulans* 04 females and males on noni or control diet (*n* = 3 biological replicates). (H) Total carbohydrate content of *D. melanogaster* CS and *D. simulans* 04 females and males on noni or control diet (*n* = 3 biological replicates). Data in (A–B), (D) and (G–H) were analyzed using two-way ANOVA, with diet and sex as factors, followed by Šídák’s multiple-comparisons test to compare diets within each sex.

**Figure S7.**
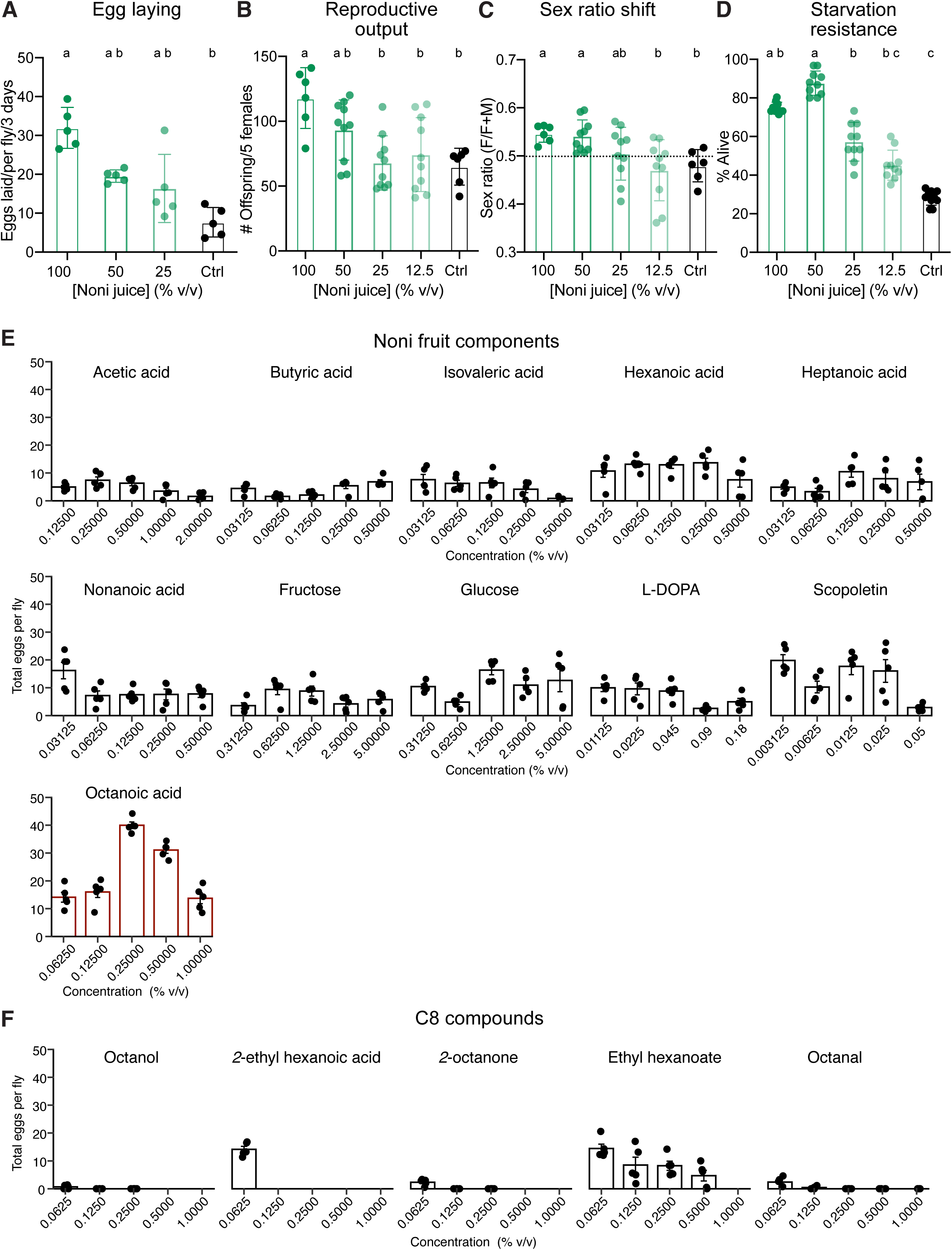
Dissection of chemical components of noni. (A-D) Dose-dependent effect of noni juice concentration on (A) egg-laying, (B) number of offspring (reproductive output), (C) sex ratio, (D) starvation resistance of *D. sechellia* 07 (*n* ≥ 5). (E) Egg-laying response of female *D. sechellia* 07 to 11 noni-associated components, each tested at five concentrations in control diet (*n* ≥ 5). (F) Egg-laying response of female *D. sechellia* 07 to octanoic acid related eight carbon compounds, each tested at five concentrations in control diet (*n* ≥ 5). Data in (A-D) were analyzed using one-way ANOVA, followed by Tukey’s multiple-comparisons test.

**Figure S8.**
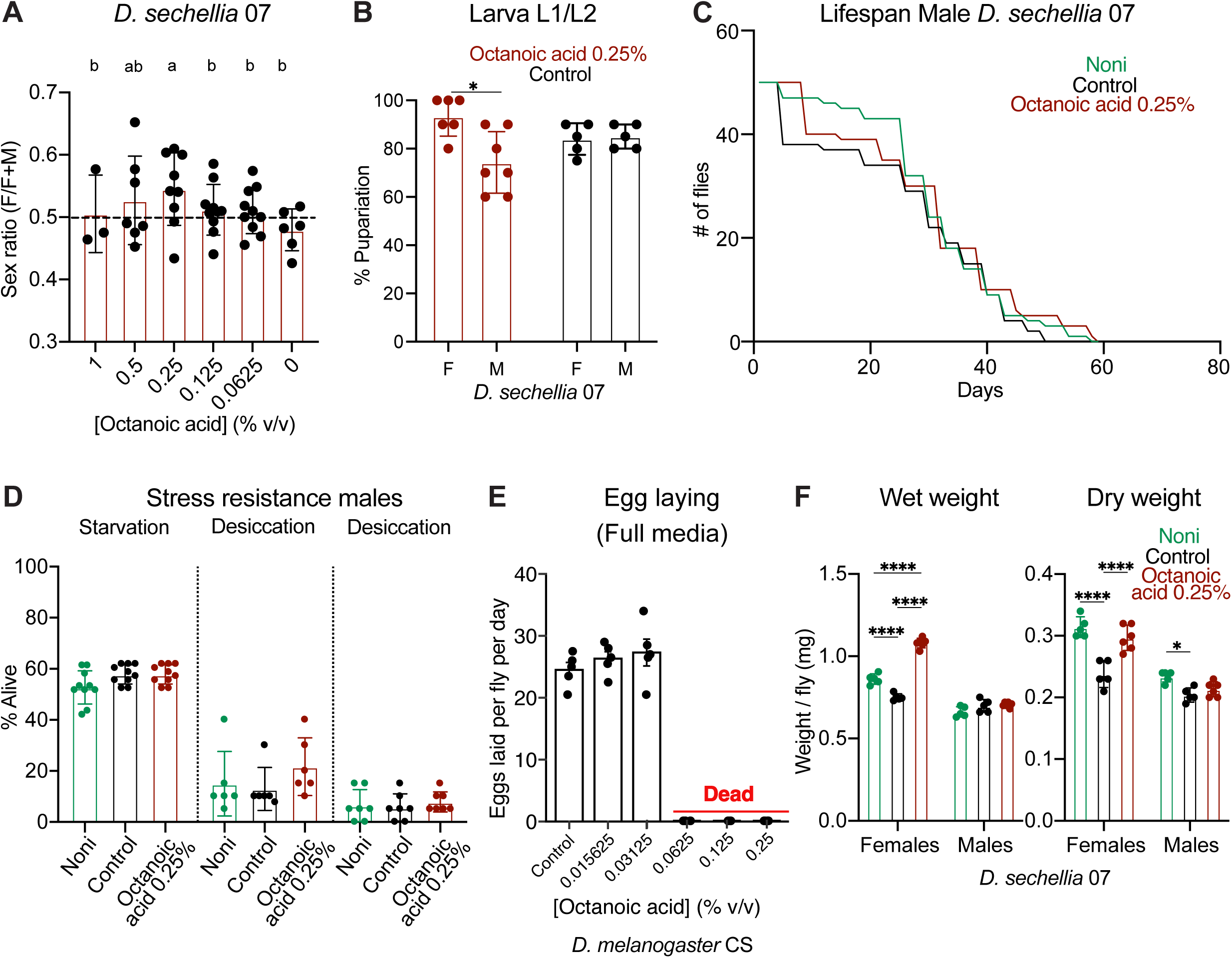
Influence of octanoic acid diet. (A) Offspring sex ratio across range of concentration of octanoic acid (*n* ≥ 5). (B) Percentage of adults emerging from L1/2 larvae placed in octanoic acid or control diet (*n* ≥ 5). (C) Lifespan of males *D. sechellia* 07 maintained on noni, control or octanoic acid diet (0.25% v/v) diets (*n* = 50 flies). (D) Stress resistance of *D. sechellia* 07 males maintained on noni, control or octanoic (0.25% v/v) diet. Starvation, desiccation, oxidative stress following 18.5 mM paraquat exposure (*n* ≥ 6 replicates). (E) Egg-laying response of female *D. melanogaster* CS to increasing concentration of octanoic acid in control diet (*n* ≥ 5). At 0.25% (v/v) octanoic acid, neither males nor females (of both *D. melanogaster* CS and *D. simulans* 04) recovered after transfer to the octanoic acid-containing diet. At concentrations above 0.0625% (v/v), all flies died without laying eggs. Control and lowest two concentration of octanoic acid diet were analyzed using one-way ANOVA, followed by Tukey’s multiple-comparisons test. (F) Wet (left) and dry (right) body weight of *D. sechellia* 07 females and males on noni, control or 0.25% octanoic acid diet (*n* ≥ 5 biological replicates of 10 flies each). Data in (A) and (D) were analyzed using one-way ANOVA, followed by Tukey’s multiple-comparisons test. Data in (B) were analyzed using two-way ANOVA, with diet and sex as factors, followed by Šídák’s multiple-comparisons test to compare sex within each diet. Survival differences in (C) were assessed using the log-rank (Mantel–Cox) test. Data in (F) were analyzed using two-way ANOVA, with diet and sex as factors, followed by Tukey’s multiple-comparisons test to compare diet within each sex.

**Figure S9.**
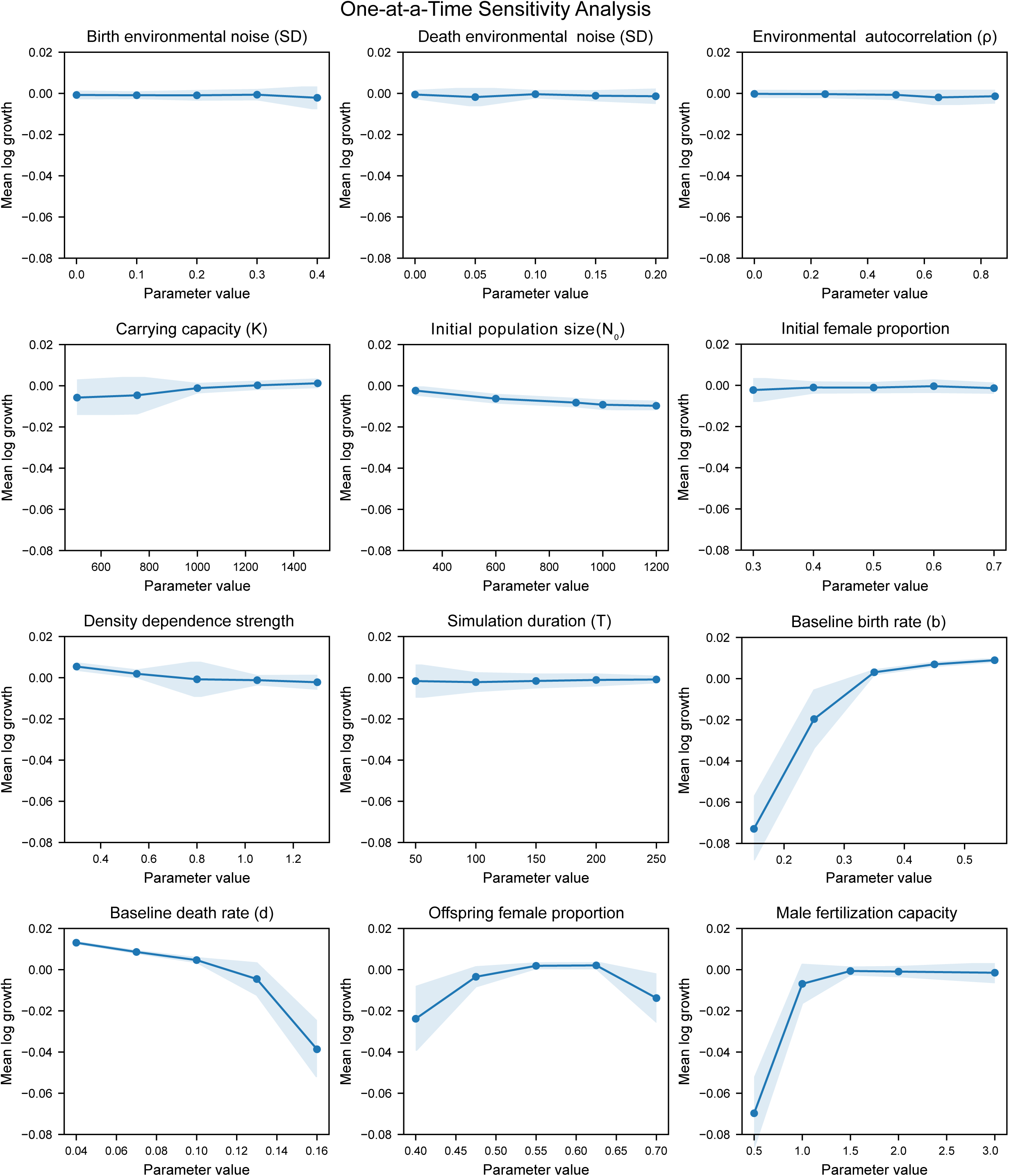
Sensitivity analysis of population dynamics to model parameters. One-at-a-time sensitivity analysis showing the effects of individual model parameters on mean log population growth per time step. Each parameter was varied across five values while all other parameters were held at their baseline settings. Each point represents the mean across 40 stochastic simulation replicates. Shaded regions indicate ±1 standard deviation across replicates. Tested parameters include environmental variability and autocorrelation, carrying capacity (*K*), initial population size (N_0_), initial female fraction, density dependence, simulation duration, birth and death rates, offspring sex ratio and male fertilization capacity.

### Supplementary Data

**Data S1.** Source data for Figure 1

**Data S2.** Source data for Figure 2

**Data S3.** Source data for Figure 3

**Data S4.** Source data for Metabolomics

**Data S5.** Source data for Figure 4

**Data S6**. Reproductive output of *D. sechellia* for different noni components

**Data S7.** Source data for Figure 5

**Data S8.** Source data for Figure 6

## Notes

### Competing Interest Statement

The authors have declared no competing interest.

